# Spatiotemporal 4D Whole-cell Modeling of a Minimal Autotroph Reveals Central Carbon Metabolism Regulated Locally by Protein Megacomplexes via Post-translational Modifications under Light Disturbance

**DOI:** 10.64898/2026.01.21.700212

**Authors:** Connah G. M. Johnson, Aaron Chan, Jordan Rozum, August George, Amar D. Parvate, Doo Nam Kim, Song Feng, Pavlo Bohutskyi, Zachary Johnson, Natalie Sadler, Marci Garcia, Xiaolu Li, Jesse Trejo, Ruonan Wu, William Sineath, Lindsey N. Anderson, James E. Evans, Angad P. Mehta, Wei-Jun Qian, Zaida Luthey-Schulten, Margaret S. Cheung

## Abstract

Photosynthetic microorganisms rely on multiple pathways in central carbon metabolism to adapt to fluctuating light and energy availability across diel cycles. Mechanistic insight into the regulatory dynamics of this adaptation requires integrating processes spanning disparate timescales, from rapid redox-dependent post-translational modifications (PTMs) to slower changes in protein expression and metabolic pathway usage. To address this complexity beyond genome-based inference and traditional modeling, we develop a whole-cell four-dimensional (3D + time) model of the marine cyanobacterium *Prochlorococcus marinus* MED4 that explicitly represents the spatial organization of enzymatic and molecular processes in central carbon metabolism under light perturbation. We employ a perturbation-based research design to experimentally generate time-series, multi-omics measurements that provide molecular descriptors and cryo-ET images as constraints for this dynamic 4D framework. The integration of experiments and modeling across defined light regimes enables quantitative validation of system-level responses and forecasting under distinct light disturbances. We test the hypothesis that light-dependent redox PTMs regulating the structural assembly of a protein megacomplex, the “dark complex,” modulate metabolic flux at a conserved regulatory node of the Calvin–Benson cycle (CBC) in cyanobacteria. Our model shows that subcellular spatial organization buffers rapid light-induced changes in thylakoid reaction rates, which are followed by redox-PTM-mediated sequestration or release of CBC enzymes in the dark complex, ultimately impacting carbon fixation dynamics within carboxysomes. Comparison with an equivalently parameterized well-mixed stochastic model demonstrates that post-translational regulation not only buffers transcriptional noise and diffusion-driven fluctuations but also stabilizes phenotypic outcomes, underscoring the importance of spatial heterogeneity in phenotypic robustness. This ability to probe adaptive, spatiotemporally resolved mechanisms in photosynthetic machinery and central carbon metabolism addresses a critical gap in genotype-to-phenotype inference and expands modeling and design capabilities for understudied or genetically intractable autotrophs such as *P. marinus* MED4.

**Significance Statement:** This work advances 4D whole-cell modeling by presenting the first spatiotemporal simulation of a photosynthetic autotroph using the Lattice Microbes platform. Using *Prochlorococcus marinus* MED4, we show that subcellular spatial organization of organelles, diffusion constraints, and redox regulation collectively shape central carbon metabolism across orders of magnitude in space and time. Through a perturbation-based strategy that generates multi-omics data sets over time, we construct and validate a spatially and temporally resolved model of MED4, constrained by high-resolution (10 nm) cryo-electron tomography. Our results highlight the importance of localized biochemical reactions and redox-dependent post-translational modification of enzymes in regulating carbon fixation in a noisy environment under light disturbance. This study establishes a spatiotemporal, whole-cell physiology modeling framework as a transformative tool for uncovering multiscale regulatory responses to environmental gradients.

## Introduction

Photosynthetic microorganisms, including model cyanobacteria, have been engineered to produce a variety of biofuels and high-value chemicals. Despite substantial progress in developing photosynthetic bioproduction platforms, insufficient and variable yields currently limit opportunities to scale up for industrial deployment (1). A central barrier contributing to its phenotypic instability is the difficulty of predicting how metabolic fluxes and regulatory proteins respond to environmental changes; cells must continuously balance energy capture from photosynthesis, carbon fixation, and redox homeostasis over a diel cycle (2) (**Figure 1A**). Photosynthetic microorganisms rely on multiple interconnected pathways in central carbon metabolism (3) to cope with light and energy limitations, principally the Calvin–Benson cycle (CBC) (4), glycolysis/gluconeogenesis, the oxidative and reductive pentose phosphate pathways (PPP), and organic carbon storage and mobilization (5).

**Figure 1.**
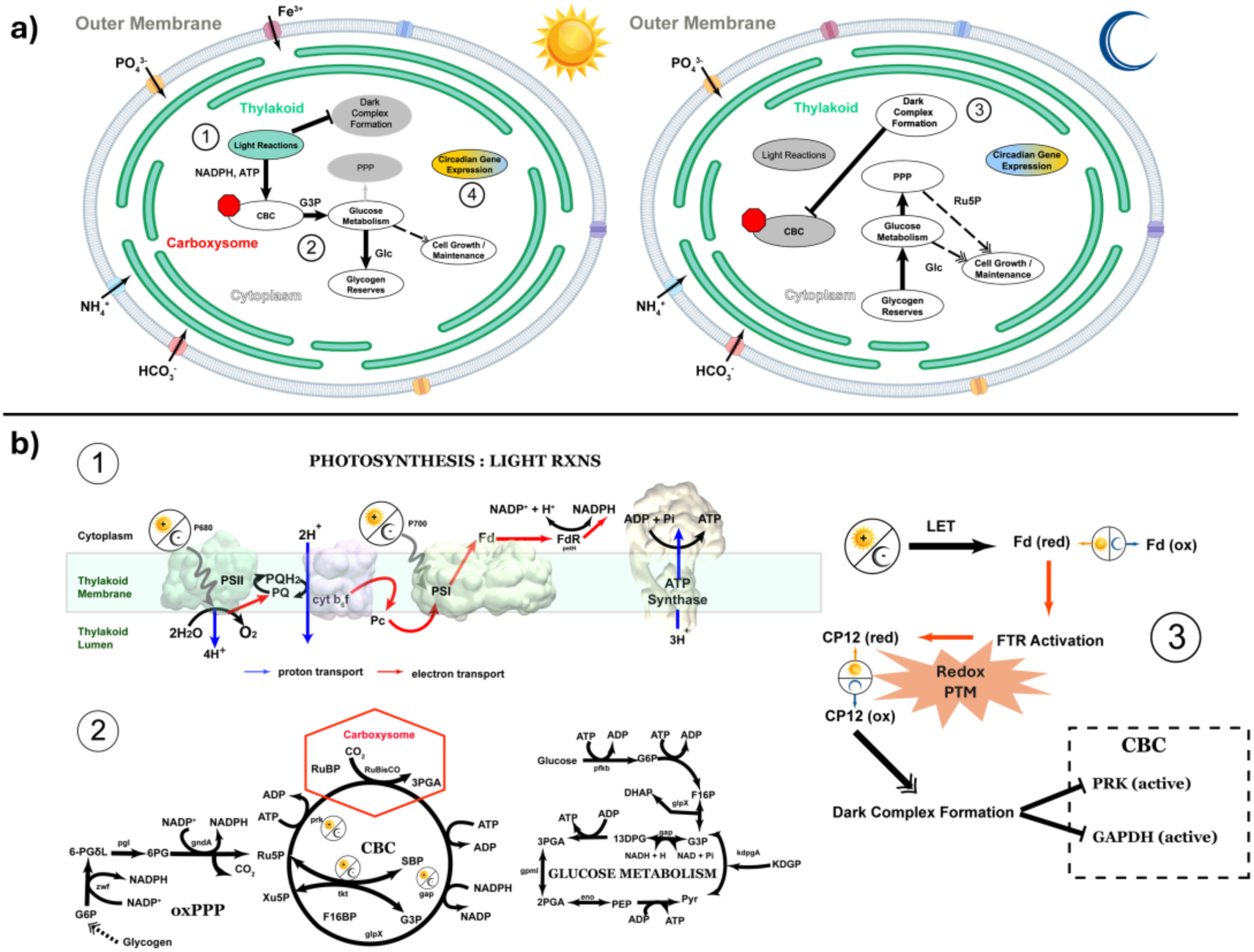
Key metabolic phenotypes in the photoautotroph MED4. The metabolic state of MED4 responds to changes in light conditions via direct energy balance, post-translational modification, and circadian entrainment. a) The influence of light exposure during the day (left) links regulatory and metabolic processes across cellular compartments. b) Detailed view of reactions involved in major energy-related metabolic subsystems including the light reactions (b.1) and the dark reactions and glucose metabolism (b.2). Redox dependent PTM of CP12 modulates inhibition of key CBC metabolic active enzymes (dimer Prk2 and tetramer Gap4) through the formation of a protein megacomplex, the dark complex (inactive Gap8CP124Prk8) (b.3). Light-powered reactions in the thylakoid (b.1) produce NADPH and ATP, which fuel carbon fixation in the carboxysomes via the CBC feeding into glucose metabolism (b.2) and ultimately glycogen storage and cell growth. In the absence of light, the oxidized format of the dark complex inhibits key CBC enzymes by sequestering them in the cytosol (b.3), and metabolic shunts favor the oxidative pentose phosphate pathway over the CBC. In this light-energy limited scenario, the cell ultimately satisfies its energetic needs (i.e. reductant NADPH) through the breakdown of its glycogen reserves.

Together with the genetic and post-translational mechanisms that modulate their activity, these pathways enable cyanobacteria to adapt to fluctuations in light and energy availability across diel cycles (6, 7). Reductant flux through these pathways must be continually rebalanced to vary with the output of the photosynthetic electron transport chain (**Figure 1B top**) with the demands of carbon fixation, biomass synthesis, and redox homeostasis during the day and oxidative PPP during the night (8) (**Figure 1B bottom**). Biodesigns that neglect the principles of regulation and redox-dependent control in cyanobacteria often fail to scale valuable bioproducts across conditions.

The primary challenge in systems modeling for phenotypic prediction of the metabolic state is capturing the vast possibilities emerging from multiple connected pathways and protein regulators at different timescales (9). One mechanism for protein regulation that is particularly relevant in photosynthetic organisms is the redox post-translational modifications (PTM) on protein cysteine residues (10) in response to changes in photosynthetic photon flux and other environmental stimuli. Nucleophilic cysteine thiols in proteins can switch reversibly between reduced and oxidized states via redox PTMs(10). This process, often known as “redox switch” (11), dynamically impacts protein conformation (7) and activity (7), thereby modulating protein-protein interactions and subsequently reconfiguring metabolic networks (12). Using illumination as a perturbation, our team previously deployed a physics-informed machine learning approach that integrated data-driven learning with correlative analysis of proteomic and transcriptomic datasets to understand complex cellular responses under light disturbance (13). On fast timescales, redox PTMs alter protein structures and interactions, facilitating the transport of electron-energy from photo-induced reactions to redox reactions in the central carbon metabolism by coupling to the circadian clock (8). On slower timescales, transcriptional and translational control, implement longer-term metabolic reprogramming (12). While capturing the regulatory layers that interact to shape phenotypes in response to environmental gradients over the entire diel cycle remains a major challenge, the use of environmental stimuli such as light as perturbations that drive the dynamics of phenotypic expression through molecular signatures has emerged as an effective strategy to integrate multimodal data into representative models that generating mechanistic insight for design and intervention.

In this study, we focus on photosynthetic light-harvesting and carbon-fixation in cyanobacteria under light disturbance at key time points in a diel cycle. A central hypothesis is that understanding the structural knowledge of molecular interactions and their dynamics with spatial resolution in response to light disruption, will provide mechanistic understanding of the interactions that orchestrate the multiple pathways in a metabolic state over a diel cycle, enabling strategies for intervention.

To test this hypothesis, we focus on the redox of molecular interactions (e.g., protein complexes) and build dynamic models of phenotypic responses to light disturbance. With the representative of a key function relating to central carbon metabolism, we leveraged a 4D whole-cell kinetic model based on the Lattice Microbes (LM) (14) to integrate multi-omics on a whole-cell framework with subcellular and cellular architecture taken from cryo-electron tomograms (cryo-ET).

Lattice Microbes (LM) (15) is a GPU-accelerated whole-cell modeling computational tool designed to simulate cellular dynamics with spatial resolution. The foundation of cellular dynamics in LM is built on a framework which integrates genetic information processing and genome-scale metabolic modeling, which requires gene annotation curated by experts.

We focus on the picocyanobacterium *Prochlorococcus marinus* MED4, the smallest known free-living photosynthetic autotroph (≈0.5 µm in diameter) (16), which provides a tractable platform with a known genome for a laboratory grown strain to mitigate risks from unknowns for experimentation (17) or for integrating data for modeling (18). Our approach begins with annotating the metagenome of the lab-strain MED4 from public databases and prior studies (19) to fill gaps in functional annotation and identify instances where regulatory proteins may be mistaken for pseudogenes. Using gene essentiality studies, and flux balance analysis (FBA) applied to genome-scale metabolic models (GSMM) as foundational tools for 4D whole-cell modeling (20), we further utilized light as a perturbation tool to drive molecular interactions and trace phenotypic responses under redox PTM at cross both temporal and spatial scales with high-resolution experimentation.

The approach of perturbation with illumination requires a coordinated research design and experimentation to generate high quality, correlative datasets including multiomics and cryoET images for model integration in this study. Ours includes perturbation studies in MED4 under controlled and varied illumination over diel cycles to capture proteome and transcriptome datasets at dawn or dusk, and each has two kinds of light variations (i.e. Dark-to-Light, Light-to-Dark, Dark-to-Dark, and Light-to-Light; **Figure 2**). These systematic datasets under illumination perturbation reveals a mechanistic representation with causal relationships between stimuli and phenotypic -omics responses, enabling intervention insights into molecular regulation mediated by PTMs. Our spatially resolved simulations focused on the role of the dark complex (21). This dark complex serves as a conserved regulatory node in the Calvin-Benson Cycle (CBC) within cyanobacteria (**Figure 1C**). Under dark (oxidized) conditions, the oxidized form of CP12 (4) binds to phosphorubilokinase (Prk) and glyceraldehyde 3-phosphate (Gap), and assemble them into an enzymatically inactive dark complex Gap_8_CP12_4_Prk_4_ (4), effectively shutting down CBC. Under light (reduced) conditions, CP12 becomes structurally disorder, the megacomplex disassembles, and release the active dimer Prk_2_ and tetramer Gap_4_ into the cytosol, switching on CBC (22). Despite its importance, the lack of spatial understanding of molecular interactions linking photosynthesis, light-dependent reactions, to light-independent central carbon metabolism over broad time scales makes it difficult to predict cyanobacterial metabolic adaptations to light over diel cycles as this process is also complicated by the gene regulation of circadian expression (23). In this study, we emphasized the importance of redox PTMs on protein megacomplexes that enable crosstalk between redox and carbon fixation pathways, exemplified by the dark complex. With a perturbation approach, this study not only generated, but also integrated correlative multiomics with causal relations, cellular imaging, reaction diffusion models, and genome scale metabolic modeling to improve mechanistic understanding of the processes that regulate photosynthetic machinery and central carbon metabolism.

**Figure 2.**
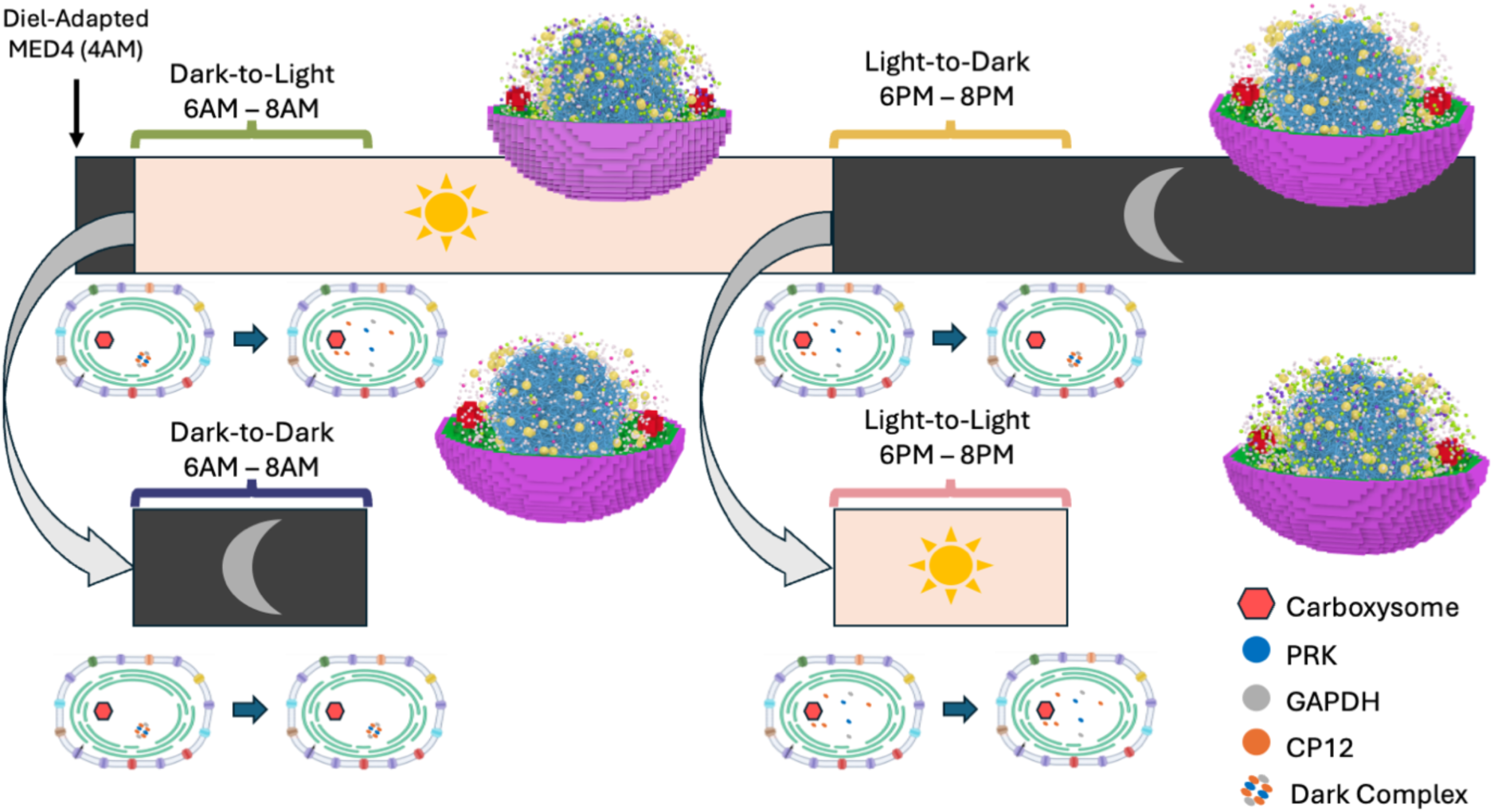
Overall research design with a perturbation approach using illumination. We adapted the cultures of MED4 to a 24-hour diel cycle consisting of 12 hours of light (6AM-6PM) alternating with 12 hours of darkness (6PM-6AM). We collect three samples each for bulk proteomics and transcriptomics in two-hour intervals over a 24-hour period starting at 6AM (dawn, Dark-to-Light scenario, just before illumination) or at 6 PM (dusk, Light-to-Dark scenario, just before dark). We consider two light perturbations in the form circadian disruption: continued darkness from 6AM-8AM (the Dark-to-Dark scenario) and continued light from 6PM-8PM (the Light-to-Light scenario). For these disrupted scenarios, we collected bulk transcriptomics data. For our Lattice Microbes simulations, we captured these four scenarios for MED4.

## Methods

### I. *Prochlorococcus* as a model photoautotroph for whole-cell simulation

We selected the picocyanobacterium *Prochlorococcus marinus* strain MED4 (CCMP 1986) as our model organism for whole-cell simulation. MED4 cells are roughly spherical and approximately 0.6 − 0.7 𝜇𝑚 in diameter (24) sufficiently thin to obviate laborious sample preparation techniques (e.g., cryo-focused ion beam milling and lamella generation) and to reduce noise from inelastic scattering at high tilt angles. Its small size enables high-resolution whole-cell tomography while minimizing computational requirements for 3D reconstruction, analysis, and storage. In addition, MED4 possesses one of the smallest (1.66 Mbp) genomes of any known photosynthetic organism encoding approximately 1,700 genes (25) which narrows the range of proteome targets for sub-tomogram averaging and reduces model complexity through minimal metabolic redundancy and a simplified regulatory architecture. As a high-light-adapted ecotype, MED4 can be readily cultivated under controlled laboratory conditions, enabling systematic perturbation of light regimes. These factors allow us to interrogate diverse molecular interaction scenarios through timely hypothesis testing using the precise environmental perturbation tools to drive behavior of interest making MED4 an excellent model organism for whole-cell modeling in the context of cross-scale systems biology.

### II. Experimental design: diel time-course for multiomics and circadian-light decoupling for expression profiling

We applied correlative *in situ* cryo-electron tomography and multiomics approaches (transcriptomics and proteomics) to monitor shifts in cellular state due to diel cycling and light perturbations in laboratory-grown *Prochlorococcus marinus* MED4, providing the empirical foundation for whole-cell modeling.

We maintained *Prochlorococcus marinus* MED4 (CCMP1986) in Pro99 medium (26) at 22 °C under a 12 h light: 12 h dark diel cycle, with lights on at 06:00 (dawn) and off at 18:00 (dusk). Photosynthetically active radiation (PAR) was ∼45 µmol photons m⁻² s⁻¹. Cultures were grown in 1 L glass bottles sparged with air and entrained to the diel cycle for at least 180 days prior to experiments.

#### Diel time-course experiment

For the diel time-course experiment, synchronized cultures were pooled from a single large culture, distributed into 50 mL bioreactors with vent caps, and destructively sampled at 2-hour intervals over 24 hours (12 timepoints, three biological replicates per timepoint). The first samples were collected at 06:00, immediately before dawn (before illumination), representing cells at the end of the night (after 12 hours of darkness). The 18:00 samples were collected immediately before dusk (before light removal), representing cells at the end of the day (after 12 hours of light). Samples were processed and prepared for global proteomics and transcriptomics (RNA-seq) separately (Please see SI for more information).

#### Circadian-light decoupling experiment

To distinguish circadian-driven responses from direct light responses, additional transcriptomics samples were collected under light perturbation conditions (**Figure 2**). After dawn, we collected samples at 08:00 under continued darkness (Dark-to-Dark: light withheld, circadian day but no light cue). After dusk, we collected samples at 08:00 under continued light (Light-to-Light: darkness withheld, circadian night but light still on). These perturbation samples, together with the corresponding 08:00 light (Dark-to-Light) and 20:00 dark (Light-to-Dark) samples from the diel time-course, enabled us to separate transcriptional changes driven by the circadian clock from those directly induced by light or darkness (13). (Please see SI for more information).

#### Sample harvesting

At each timepoint, samples were processed separately for transcriptomics and proteomics analysis. For transcriptomics, 10 mL of culture was harvested by centrifugation (6,500 × g, 6 min, 4°C). Cell pellets were snap-frozen in liquid nitrogen and stored at −80°C until RNA extraction. For proteomics, cellular metabolism was quenched by mixing 50 mL of culture with an equal volume of 60% methanol pre-cooled to −20°C, yielding a final concentration of 30% methanol to preserve the in situ proteome. Samples were centrifuged (6,500 × g, 6 min, 4°C), and cell pellets were rinsed once with ice-cold PBS, snap-frozen in liquid nitrogen, and stored at −80°C until protein extraction.

Together, these experiments generated correlative multiomics datasets representing diel dynamics and four perturbation states, i.e., a circadian-aligned (“Dark-to-Light”, “Light-to-Dark”) or circadian-decoupled (“Dark-to-Dark”, “Light-to-Light”) light conditions across two-hour intervals, providing empirical constraints for parameterizing Lattice Microbes simulations. Detailed protocols for culture preparation and sample processing are provided in Supplementary Information (Sections **S1 – S4**).

### II.2 GENERATION OF MULTI-OMICS DATA

#### Genome Re-annotation and Functional Analysis

##### Reference genome acquisition and re-annotation

The *Prochlorococcus marinus* MED4 (GCA_000011465.1) was obtained from NCBI together with the corresponding GFF and protein FASTA files. To ensure completeness of protein-coding gene sets, genes were predicted de novo from the reference genome using Prodigal (v2.6.3) (27), translated into protein sequences, and reconciled with NCBI-provided gene models to identify missing, truncated, or alternatively annotated genes. This reconciliation step mitigates known annotation biases, particularly those affecting short regulatory proteins such as CP12, which are frequently annotated as pseudogenes or omitted from public records. Following coordinate validation, proteins were deduplicated by genomic position.

##### Functional annotation

Reconciled protein sequences were functionally annotated using complementary approaches to maximize coverage across both conserved and divergent proteins. KEGG Orthology assignments were generated using KOfamScan (exec_annotation, v1.3.0) against curated KOfam HMM profiles with default score thresholds. Conserved domain annotation was performed against the Pfam-A database (release 2024-10-15) using HMMER hmmscan (v3.3.2) (28) with trusted cutoffs (--cut_tc). Additionally, enzyme function predictions were generated using CLEAN (v1.0.1) (29), a deep learning-based framework that produces per-protein maximum separation scores and associated p-values. Annotation outputs were integrated using a custom Python-based consensus pipeline that reconciled protein identifiers across sources, taking the majority rule to determine the Enzyme Commission (EC) number from curated high-confidence homology-based annotations (KEGG Orthology and Pfam) and CLEAN-derived predictions while retaining method-specific evidence for transparency. For proteins assigned KEGG Orthology identifiers or EC numbers, associated KEGG reaction identifiers were retrieved via the KEGGREST API to enable pathway-level interpretation. Curated auxiliary metabolic gene reference mappings were incorporated based on high-identity UniProt matches (>95% identity; database accessed 2024-08-28). Basic protein properties, including hydrophobicity profiles and predicted membrane association, were calculated and are included in the annotation table provided with the data release (30).

#### Transcriptomics

##### RNA extraction and sequencing

Total RNA was isolated from three independent biological replicates per time point using the RNeasy kit combined with QiaShredder homogenization columns (QIAGEN). Briefly, cells were lysed in RLT buffer supplemented with 1% β-mercaptoethanol and passed through QiaShredder columns for homogenization. Following addition of 70% ethanol, samples were processed according to the manufacturer’s protocol. Each biological replicate was carried through library preparation and sequencing independently. Prior to library construction, samples underwent DNase treatment and ribosomal RNA depletion. RNA integrity was verified by the sequencing facility before proceeding with library preparation. Paired-end sequencing (2×150 bp) was performed on an Illumina platform by Azenta US, Inc., yielding approximately 350 million reads (∼105 GB) per sample with single indexing.

##### RNA-seq Data Processing and mRNA quantification

Sequencing reads were mapped to the *Prochlorococcus* MED4 reference genome. Adapter trimming and quality filtering were performed using TrimGalore (v0.6.10) (31) with CutAdapt (v5.0) (32), and read quality was evaluated with FastQC (v0.12.1) (33). Gene-level read counts were generated using Rsubread (v2.20) with default parameters (34). Alignment quality metrics were obtained through Samtools (v1.21) (35) and Qualimap (v2.3) (36), with consolidated multi-sample quality reports generated via MultiQC (v1.28) (37). Functional annotations were assigned using annotation outputs described above.

#### Global proteomics

##### Sample processing

Proteomics samples were processed by MPLEx method as described previously (38). Briefly, cell pellets were resuspended in ice-cold Milli-Q water, then lysed by resuspension in cold 2:1 chloroform:MeOH (stored at – 20°C) plus intermittent vortexing and sonication. Samples were centrifuged at 4°C, 10000g for 10min to pellet proteins. Protein pellets were collected and dried with N_2_ stream, then resuspended in 100mM ammonium bicarbonate with 8M urea (pH 8). Then, samples were reduced by 5mM dithiothreitol at 60°C for 30min, then alkylated by 20mM iodoacetamide at 37°C for 1hr. The samples were diluted by 8-fold with 100mM ammonium bicarbonate (pH 8). Protein digestion was conducted with trypsin (trypsin:protein = 1:50, w/w) and 1mM CaCl_2_ at 37°C for 3hrs. The digested peptides were desalted by SPE C18 cleanup procedure, then subjected to MS analysis.

##### LC-MS/MS analysis

Proteomics samples were analyzed by an Orbitrap Astral Mass Spectrometer coupled with a Vanquish Neo UHPLC System (Thermo Scientific, CA). An EASY-Spray HPLC column (150mm × 150µm inner diameter) was used for separation (with 14 min gradient). The data-independent acquisition (DIA) mode was used. The isolation window was 2 m/z. Precursor mass range was 380 – 980 m/z. Raw MS data were analyzed by DIA-NN (39) against the annotated protein database we curated as described in the previous session. Fully tryptic search was conducted. Cysteine carbamidomethylation (+57.0215 Da) was enabled as a fixed modification. Methionine oxidation (+15.9949 Da) and acetylation on N-terminal (+42.0106 Da) were included as dynamic modifications.

##### Proteomics data processing and iBAQ calculation

The raw MS spectral data are processed using DIA-NN and the resulted LFQ intensity quantification of protein abundance are used to calculate the iBAQ values using DIAgui (40). We estimated protein counts from iBAQ values by calculating the scaling factor to convert iBAQ to copy numbers for which a target total protein mass is obtained. This target mass is computed from the median cell mass of 66fg (41), and the protein dry mass fraction of 58% reported by (42, 43).

### III. Whole-autotroph bioimage capture

*Prochlorococcus marinus* MED4 (44, 45) cells from diel-entrained cultures collected during the light period were concentrated to (∼10^9^ cells/ml) for preparing samples for cryo-electron tomography (Cryo-ET). Briefly, 3 µL suspension of MED4 cells (∼10^9^ cells/ml) mixed with mixed with 10 nm BSA coated colloidal gold fiducials (Aurion SKU: 200.133) was loaded on to holey carbon grids (Quantifoil Q2/2 or 2/1, 200 or 300 mesh). Grids were vitrified by plunge freezing in liquid ethane using a Leica EM GP2. Gird screening and data collection were performed on the 300 keV TFS Titan Krios G3i cryo-electron microscope at the Environmental Molecular Sciences Laboratory (EMSL), Pacific Northwest National Laboratory (PNNL). Tomographic data were collected on a Gatan K3 (Gatan Inc.) camera using a Bioquantum energy filter, with the slit width at 20 e^-^V, using a Volta Phase Plate, at 33,000 x nominal magnification. Tilt series were collected from -/+54^0^ at 3^0^ intervals using SerialEM (44). Individual tilt images were recorded as movies with 10-12 subframes and an exposure of 1.02 s in super resolution mode resulting in pixel size of 1.3 Å and a total dose of 110-120 e^-^/Å^2^ and a nominal defocus of -4 µm. Custom in-house scripts were used to run the automated workflow of image processing from motion correction to final 3D tomogram calculation. MotionCorr2 (https://doi.org/10.1038/nmeth.4193) was used to perform motion correction of the individual movies. Restacking and CTF correction and reconstruction of the tomograms using IMOD (https://doi.org/10.1006/jsbi.1996.0013) and ETomo suite (https://doi.org/10.1016/j.jsb.2016.07.011). Final reconstructed tomograms were binned 8x with a pixel size of 10.4 Å relative to the raw data and visualization of the 3D volumes was done using 3dmod. The tomogram was segmented using MemBrain-seg (46, 47), ColabSeg (48), and ChimeraX (49) with ribosome particle locations picked using crYOLO (50).

### IV. Flux Balance Analysis (FBA)

We incorporate the genome-scale metabolic model (GSMM) of MED4 developed by (43), which builds upon the work of (42), into our model to represent the metabolism. The metabolic component of the spatiotemporal model is evaluated using the flux balance analysis (FBA) approach for which it was designed (43). This model integrates 994 reactions among 802 metabolites governed by 595 metabolic genes via curated gene rules that couple gene expression to reaction activity, **Figure S1**. In addition to core metabolic reactions, the model incorporates nutrient uptake and usage, cell maintenance, biomass formation (including nucleotide and amino acid synthesis, see **Figure S2**), and glycogen storage and usage as shown in **Figure 3**. When simulating the model, we incorporate enzyme usage constraints derived in real-time from enzyme kinetic parameters and the spatial part of the simulation. In this way, we couple spatially resolved protein abundance and PTM information from the reaction diffusion component of the model to metabolic fluxes. We then optimize biomass accumulation subject to these enzymatic constraints, cell maintenance constraints, photosynthetic photon input, and glycogen storage or usage. We then perform a second optimization pass using a parsimony objective (minimization of total absolute flux), which is a standard approach, called parsimonious FBA (pFBA), for reducing degrees of freedom in GSMMs (51, 52). In all cases for the FBA, we perform a multi-step optimization process that prioritizes maintaining cell function (modeled as flux through a *maintenance* reaction), followed by biomass accumulation, and lastly glycogen storage if applicable. Because we explicitly model the effect of the molecular dark complex in regulating CBC, we enforce the usage of free (non-sequestered) enzyme.

**Figure 3.**
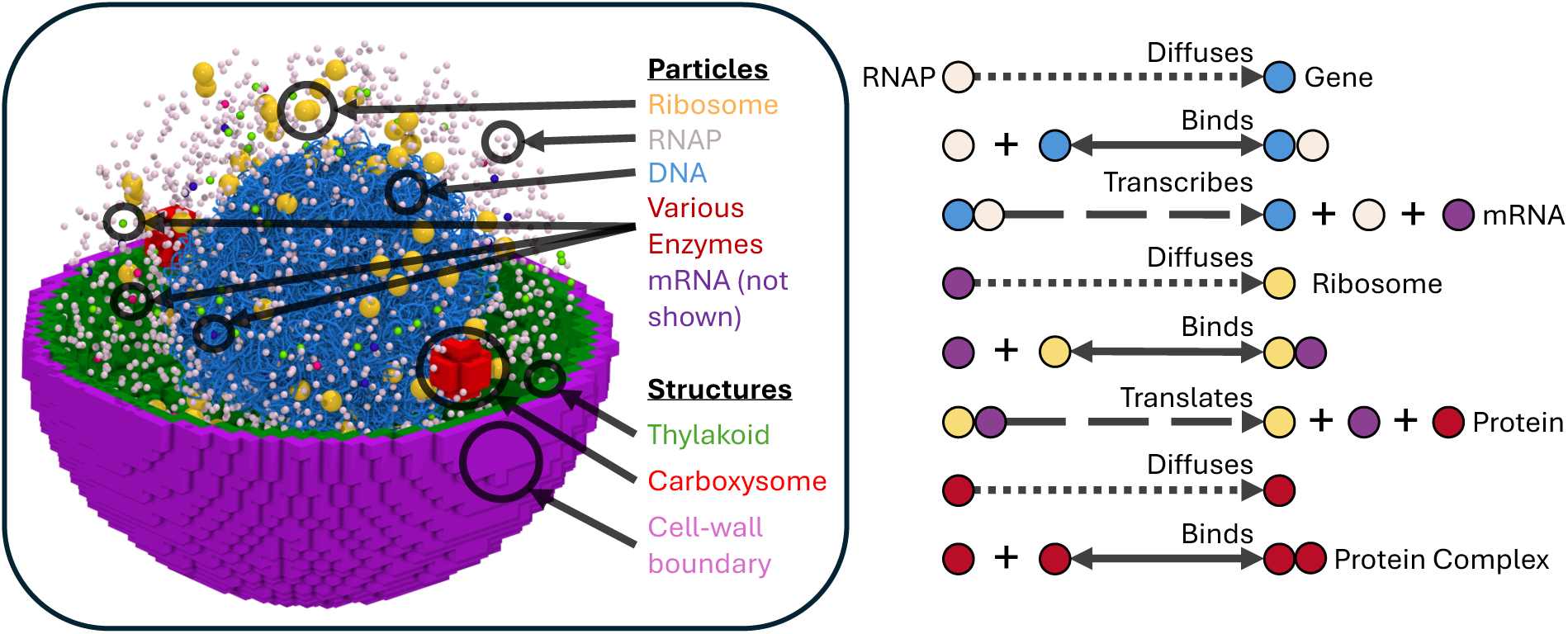
Overview of the reaction diffusion portion of the model from Lattice Microbes. The spatially heterogeneous simulation domain is depicted on the left, with key diffusing particles and static cellular substructures highlighted. A schematic of the reaction and diffusion processes incorporated into the model is depicted on the right. Degradation and dissociation reactions are omitted from the schematic. The depicted processes shown are implemented and separately parameterized for each spatially resolved protein.

### V. Lattice Microbes: whole cell 4D kinetic modeling framework

The whole-cell model for *Prochlorococcus marinus* MED4 was constructed using Lattice Microbes (LM) 2.4 [https://github.com/Luthey-Schulten-Lab/Lattice_Microbes/releases/tag/v2.4] leveraging the reaction-diffusion master equation (RDME) to capture spatially and temporally resolved molecular interactions throughout the cell (**Figure 3**). The simulation volume is subdivided into cubical voxels, and each voxel is assigned a site type (e.g., cytosol, extracellular medium, thylakoid, carboxysome). Large macromolecular complexes such as ribosome, carboxysomes, and DNA are represented as voxels or groups of voxels. Smaller species, such as individual proteins or transcripts, are modeled as particles that stochastically diffuse between and react within voxels. Our model uses a spatial resolution of 20 nm, with a temporal resolution of 8 µs to ensure that the root mean squared displacement per time step of each diffusing object tracked in the simulation remained below the resolution threshold. The spatial resolution was selected to be as coarse as possible (for computational efficiency) while preserving the distinction between the cell wall and the thylakoid. The temporal resolution was selected such that the root mean squared displacement of enzymes within a single time step did not exceed the voxel width (see Methods Section V.1 for further details regarding enzyme diffusion parameters).

The two initial conditions (dawn or dusk) of these simulations were parameterized from the -omics experiments described in Method Section II (see Methods Section V.3 for details). To systematically explore physiological behavior to incorporate light on or off, we simulated the cell’s response at dawn or dusk to either a continuation (*Dark-to-Dark*, *Light-to-Light*) or abrupt change (*Dark-to-Light*, *Light-to-Dark*) in light intensity over the course of two hours. We used LM’s in-built support for coupling to other models to integrate the FBA-derived fluxes of the GSMM described in the previous subsection to quantify biomass production rates and estimate metabolic activity under various light and dark conditions (**Figure 4**). The transcription and translation of spatially organized genes, and degradation of the resulting mRNA and protein molecules were defined by specifying appropriate reactions and diffusion parameters. These parameter sets were derived through experimental investigation in this study, literature/database searches as described in the following subsections or estimated through computational packages such as AlphaFold2 (53). We provided a schematic overview of the workflow with input-output relationships between data and modules for Lattice Microbes software when it updates a cell state in **Figure 4**.

**Figure 4.**
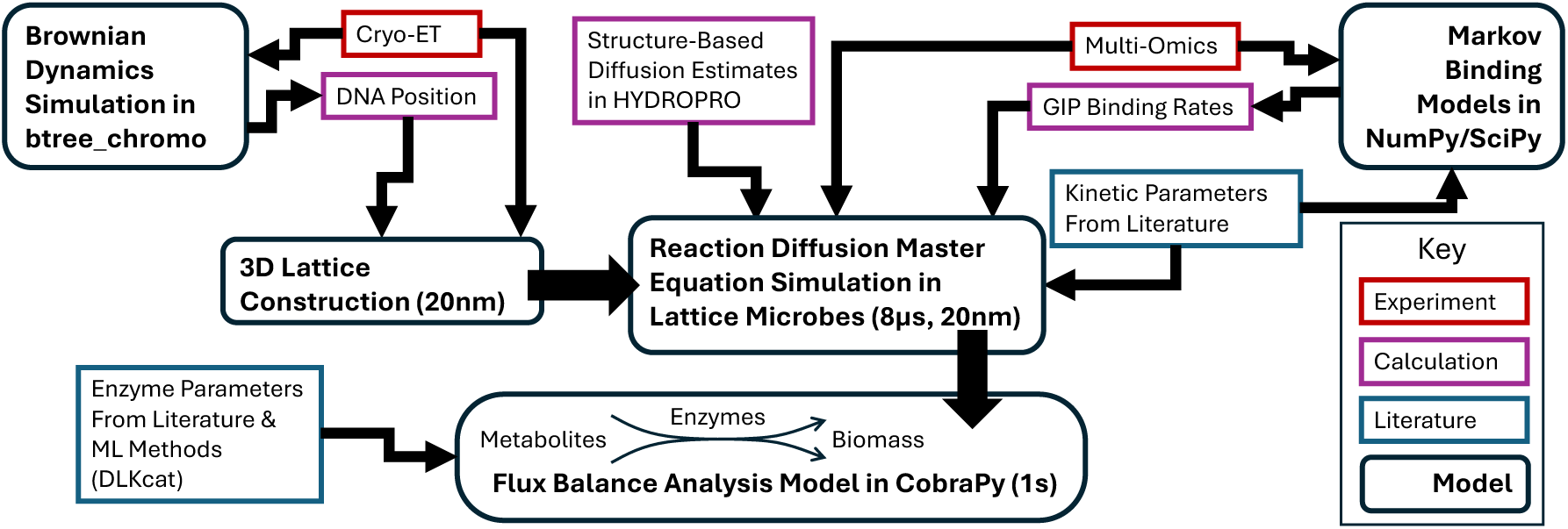
The modules of the Lattice Microbes software combine to update the cell state. Small arrows indicate input-output relationships between data and modules. Large arrows indicate the main simulation process. Boxes with rounded corners represent model modules, while rectangular boxes indicate data. The color of each rectangular box indicates the source of the data (measured experimentally by the authors for this work, computed from simulation outputs, or gathered from the literature for red, purple, and blue, respectively).

### V.1 Parameterizing the 3D spatial model

Cryo-ET data provided high-resolution visualizations of critical subcellular structures such as cell wall, cell membrane, thylakoid, carboxysomes, and ribosomes. We rasterized 20 nm resolution 3D lattice regions for Lattice Microbes following the method (54) to approximate the shape of the segmented masks from cryo-ET data. This procedure maps the simulation domain onto a 64 x 64 x 64 3D lattice with a grid spacing of 20 nm. The discretized cell is an ellipsoid with widths 32 x 28 x 28 voxels (720 nm x 560 nm x 560 nm). Because the picked ribomsomes represented a small fraction of its protein abundance, we therefore placed the ribosomes randomly within the cytoplasm region and they were prevented from being placed within 1 voxel of the carboxysomes or thylakoid regions.

Genomic data, coupled with Brownian Dynamics (BD) simulations (e.g., btree_chromo (55)), established the spatial positioning of genes encoding CBC enzymes throughout the circular genome. The circular genome of MED4 was weaved into the cytoplasm region using gen_sc (55). We used the packed genome configuration as an input to Brownian dynamics (BD) simulation, which simulates chromosome dynamics, using btree_chromo (55) and LAMMPS (56) in order to initialize gene placement within the cell. This method models the DNA as a cyclic polymer where each monomer represents a 10 bp (3.4 nm) long DNA segment with excluded volumes. The spatial configuration of the chromosome is modelled using a BD simulation that uses a Weeks-Chandler-Andersen (WCA) potential in order to avoid self-overlap and steric collision with ribosomes and carboxysomes (57). The gene positions from the end of the BD simulation were mapped onto simulation lattice described above. Lastly, the knowledge about the proteins, RNAP, and mRNA from the correlative experiments in this study were initialized at available lattice sites in the cytoplasm. An ensemble of these stochastic position solutions was formed, resulting in different spatial configurations for the genes that encode proteins used in the metabolic model.

Regarding the diffusion coefficients, they were calculated using the HYDROPRO software (58); a correction factor was applied to account for reduced effective diffusion rates, which arise from crowding interactions in prokaryotic cytoplasms (59). **Figure 5** shows the workflow of the data transformation from captured bioimages (a) from cryo-ET described in Method II, and segmentation results with ribosome picks (b), to the 3D spatial model with inserted localized protein complexes and a simulated DNA for Lattice Microbes simulations (c).

**Figure 5.**
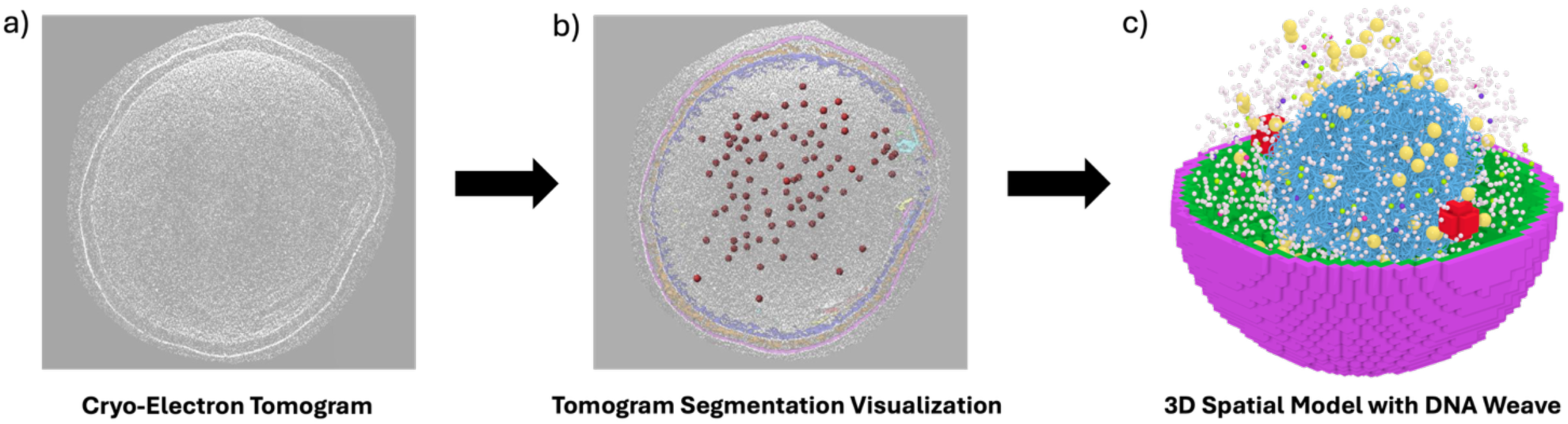
Process workflow for developing the spatial map from cryo-electron tomogram from Prochlorococcus marinus MED4. The tomogram (a) is segmented per internal, structural compartmentalization, which are shown on the tomogram along with ribosome particle picks (b). These masks are approximated as 3D regions and combined with the localized protein complexes and DNA weave to initialize the simulation (c).

### V.2 Gene selection and the gene products

We have focused the spatial component of our simulation on the primary energy metabolism of MED4, with special emphasis on how carbon fixation is regulated by changes in light conditions. Our simulation models the expression for a subset of 15 genes within the MED4 genome (**Table 1**). They topological analysis of the model and from the literature as shown in the table. We also validate them with our experimental transcriptome data (see Data Package (30)).

**Table 1.** Fifteen spatially tracked proteins selected for their regulatory or catalytic importance in the dark-light transition and carbon metabolism in MED4.

| Gene Locus | Common Name | Description |
| --- | --- | --- |
| PMM0023 | *gap2* | Enzyme in CBC; forms dark complex with Prk and CP12 |
| PMM0128 | *rpaA* | Master transcriptional regulator and readout of the kaiC circadian protein oscillator |
| PMM0208 | *eno* | Enolase, key enzyme in glycolysis |
| PMM0225 | *kdgA* | Catalyzes pyruvate synthesis |
| PMM0246 | *ntcA* | Key regulator of nitrogen metabolism |
| PMM0706 | *phoR* | Key regulator of phosphate metabolism |
| PMM0767 | *glpX* | Enzyme in CBC |
| PMM0770 | *gndA* | Enzyme in CBC |
| PMM0785 | *prk/cbb* | Enzyme in CBC; forms dark complex with Gap and CP12 |
| PMM1075 | *petH* | Oxidizes Ferredoxin in the thylakoid |
| PMM1150 | *trxB* | Global redox response protein |
| PMM1434 | *gpmI* | Enzyme in glycolysis |
| PMM1509 | *fusA* | Elongation factor |
| PMM1619 | *rer4* | Histidine kinase-modulated response regulator involved in high-light response |
| PMM1627 | *pfkB* family<br>carbohydrate kinase | Fructokinase |

These enzymes were selected from the giant connected component of the metabolic subnetwork formed from the photosynthesis, carbon fixation, and glycolysis/gluconeogenesis pathways. These genes correspond to cytoplasmic enzymes from key metabolic pathways in central metabolism, transcription factors which modulate nutrient uptake (43) or have shown notable changes expression profile under varying lighting conditions (16). We further narrow our selection based on enzyme parameters obtained as described in section V.3 of Methods; we focus on slowly diffusing (slower than 100 µm^2^/s in water), limiting enzymes with turnover numbers below 500/s and estimated protein counts below 2000. The spatially tracked gene products, enzymes, and their catalytic reactions are given in **Table 2**. The intermediate complexes containing Gap, Prk, and CP12 that lead to the formation of the dark complex are summarized in **Table 3**.

**Table 2:** The spatially tracked enzymes represented in voxels from the selected genes in Table 1 and their catalytic reactions.

| Gene Name<br>(NCBI ID) | KEGG ID | Enzyme Name | Reaction |
| --- | --- | --- | --- |
| *gap* (PMM0023) | R01061 | Glyceraldehyde 3-phosphate dehydrogenase | D-Glyceraldehyde-3-phosphate + NAD + Orthophosphate ⇌ 3-Phospho-D-glyceroyl-phosphate + H + NADH |
| | R01063 | | D-Glyceraldehyde-3-phosphate + NADP + Orthophosphate ⇌ 3-Phospho-D-glyceroyl-phosphate + H + NADPH |
| *eno* (PMM0208) | R00658 | Enolase | 2-Phospho-D-glycerate ⇌ H<sub>2</sub>O + Phosphoenolpyruvate |
| *kdgA* (PMM0225) | R05605 | 2-keto-3-deoxy-6-phosphogluconate aldolase | 2-Dehydro-3-deoxy-6-phospho-D-gluconate ⇌ D-Glyceraldehyde-3-phosphate + Pyruvate |
| *glpX* (PMM0767) | R01845 | Fructose-1,6-bisphosphatase/sedoheptulose-1,7-bisphosphatase | H<sub>2</sub>O + Sedoheptulose-1-7-bisphosphate → Orthophosphate + Sedoheptulose-7-phosphate |
| | R01070 | | D-Fructose-1-6-bisphosphate ⇌ D-Glyceraldehyde-3-phosphate + Glycerone-phosphate |
| | R04780 | | D-Fructose-1-6-bisphosphate + H<sub>2</sub>O → D-Fructose-6-phosphate + Orthophosphate |
| *gndA* (PMM0770) | R10221 | 6-phosphogluconate dehydrogenase | 6-Phospho-D-gluconate + NAD ⇌ CO<sub>2</sub> + D-Ribulose-5-phosphate + H + NADH |
| | R01528 | | 6-Phospho-D-gluconate + NADP → CO<sub>2</sub> + D-Ribulose-5-phosphate + H + NADPH |
| *prk* (PMM0785) | R01523 | Phosphoribulokinase | ATP + D-Ribulose-5-phosphate ⇌ ADP + D-Ribulose-1-5-bisphosphate |
| *petH* (PMM1075) | R01195 | Ferredoxin-NADP oxidoreductase | 4 H + 2 Plastoquinone + 4 Reduced-ferredoxin → 4 Oxidized-ferredoxin + 2 Plastoquinol |
| *gpml* (PMM1434) | R01518 | Phosphoglycerate mutase | 2-Phospho-D-glycerate ⇌ 3-Phospho-D-glycerate |
| *pfkB* (PMM1627) | R00867 | Fructokinase | ATP + D-Fructose → ADP + beta-D-Fructose-6-phosphate |
| | R00299 | | ATP + D-Glucose → ADP + D-Glucose-6-phosphate |
| | R00760 | | ATP + D-Fructose ⇌ ADP + D-Fructose-6-phosphate |

**Table 3:** Spatially tracked dark complex constituents, including active forms of Gap and Prk enzymes. Every copy of each of these complexes or monomers is localized to a cubic voxel 20 nm across at each time point. Dark complex formation is initiated by oxidation of CP12, allowing it to bind to the Gap4 tetramer. Dark complex dissociation is initiated by reduction of CP12 in-complex.

| Dark Complex Components | Notes |
| --- | --- |
| **Gap** | Monomer, coded by *gap2* gene |
| **Prk** | Monomer, coded by *prk* gene |
| **Gap<sub>2</sub>** | Not active form |
| **Gap<sub>4</sub>** | Active form of Gap enzyme |
| **Prk<sub>2</sub>** | Active form of Prk enzyme |
| **Gap<sub>4</sub>CP12<sub>2</sub>** | Oxidized CP12 binds Gap<sub>4</sub>, reducing enzymatic efficiency by approximately 50% |
| **Gap<sub>4</sub>CP12<sub>2</sub>Prk<sub>2</sub>** | Neither Gap nor Prk is active in this form |
| **Gap<sub>4</sub>CP12<sub>2</sub>Prk<sub>4</sub>** | Neither Gap nor Prk is active in this form |
| **Gap<sub>8</sub>CP12<sub>4</sub>Prk<sub>2</sub>** | Neither Gap nor Prk is active in this form |
| **Gap<sub>8</sub>CP12<sub>4</sub>Prk<sub>4</sub>** | Fully formed dark complex |

In addition to these energy metabolism enzymes, we also track six genes that are involved in major regulatory processes as described in literature. Several of these are involved in nutrient uptake regulation: *ntcA* (60), *phoR* (61, 62), and *rer4* (63) are involved in regulating nitrogen, phosphate, and light usage in the cell respectively. We also track *trxB*, a key redox indicator (64, 65), and *rpaA*, the master circadian regulator in MED4 (66). Since the circadian state of MED4 is primarily determined by the phosphorylation, rather than the abundance, of the RpaA transcription factor, we also track one of its targets in MED4, *fusA*, which is a conserved bacterial elongation factor (67) transcribed when RpaA is phosphorylated (maximal activation occurs at dusk).

### V.3 Parameterization of rates in biochemical reaction and diffusivity for dynamical simulations

In our chemical master equation (CME) and reaction-diffusion master equation (RDME) simulations, we spatially track a subset of important regulatory and metabolic proteins, their complexes, and their transcripts (as described in the previous section). In the RDME simulations we initialize the spatial positions of all genes in the genome, ribosomes, the thylakoid, cell membrane, and carboxysomes according to their cellular localization suggested by cryo-ET and BD simulations. Each spatially tracked protein therefore requires several key parameters to be specified.

These include elongation rates for translation and transcription, RNAP-promoter binding rates, mRNA-ribosome binding rates, diffusion coefficients for each cellular region (though typically diffusion is permitted only in the cytosol), degradation rates for proteins and transcripts, binding and unbinding rates, and catalytic turnover numbers (for enzymes). Many of these important rates depend on the regulatory state of the cell, which we view here as a function of the light condition. Moreover, key aspects of the well-mixed metabolic component of the model, such as which reactions are permitted to carry flux, depend on the regulatory state as well.

In general, we have set initial concentrations and counts according to multiomics quantifications at a single time point (68), and we have set genetic information processing rates according to differences in quantification over a two-hour interval. We then used these, along with data from the literature (see this section for details), to parameterize initial conditions, RNAP binding rates, and protein degradation rates. We used global values for mRNA-ribosome binding affinity and maximum mRNA and protein elongation rates. Thus, the production rate of each tracked protein was predicted from the binding rate between its RNAP and the corresponding gene’s promoter region that we inferred from transcript counts (30). Some intrinsic parameters, such as diffusion coefficients or turnover numbers, are computed in a manner that is independent of the multiomics measurements or are gathered from literature. In this section, we describe the parametrization method in detail.

For macromolecular complex binding (RNAP-DNA binding and mRNA-ribosome binding), we fit rates using a simple Markovian model (see Supplementary Information Section S5 for details). We consider three states: bound (state B), unbound prior to transcription or translation completion (state U), or unbound following transcription or translation (state T). Transition rates are computed as functions of binding and unbinding rates assuming a Poisson process. The transition rate from B to U uses a fixed small unbinding rate (0.001/s). The transition rate from B to T is fixed by the maximum transcript or protein elongation rate using values measured in *E. coli* (1.5kbp/min for transcription or 10AA/s for transcription (69)). Direct transitions between states U and T are forbidden. The transition rates from state U to B and from T to B are assumed to be equal and derived from an unknown rate *b*. By finding the eigenvector of the principal eigenvector of the transition matrix, we identified the average steady state transcription or translation rate per transcript as a function of *b*.

#### Transcriptional parameters

We estimated initial conditions for mRNA counts per cell by normalizing transcripts per million values per transcriptomics sample to the previously reported median cell mass of 66fg (41), the RNA dry mass fraction of 4.7% reported by (42, 43), assuming 80% of RNA mass is rRNA (in line with values reported in literature for other organisms (70, 71)). We used previously published mRNA decay rates derived from *in vivo* measurements in MED4 (72). To estimate transcriptional efficiency of RNAP, we constructed a simple Markov binding model, as described earlier in this section, for each gene in which DNA-bound RNAP transcribes at an average rate that is sufficient to sustain the maximum observed transcript count across all experimentally measured time points for that gene (73). RNAP-gene binding rate serves as a proxy for promoter strength and was computed numerically by fitting to change in average transcript counts over the two-hour measurement interval in each light condition (30).

#### Protein translation parameters

We estimated protein counts using proteomics-derived intensity-based absolute quantification (iBAQ) normalized to a median cell mass of 66fg (41), and the protein dry mass fraction of 58% reported by (42, 43). For translation, we assume a universal mRNA-ribosome binding rate. To estimate this rate, we fit binding parameters in a simple Markov model, as described earlier in this section, in which maximal average translation rate is proportional to the average transcript count (which is of order 1 or less). To estimate a global mRNA-ribosome binding rate, we fit a simple linear growth and exponential decay function using the median transcript length (from the gene annotation) and median estimated count (from proteomics data (30)) such that we recovered a doubling time for protein abundances equal to the reported doubling time of approximately 24hrs for MED4 (74). We used the median binding rate across the entire proteome as the *de facto* mRNA-ribosome binding rate constant. Protein decay was estimated based on the assumption that the largest fractional decrease in protein count occurs when transcription is negligible. We modeled competition for ribosome by untracked mRNA by implementing a stochastic state switch for each ribosome to simulate occupation by untracked mRNA, using mean total mRNA count and mean protein length to determine switching rates.

#### Metabolic parameters

Spatially tracked enzymes were selected from the giant connected component of the metabolic subnetwork formed from the photosynthesis, carbon fixation, and glycolysis/gluconeogenesis pathways as described in Section V.2. Particularly, we focused on the representative enzymes (shown in **Table 2)** in energy metabolism that have a high propensity to be rate-limiting and spatially heterogeneous.

For these spatially tracked enzymes (shown in **Table 2**), we constrain reaction flux using enzyme turnover numbers (k_cat_) from BRENDA (75) or inferred using machine learning methods (76) wherever unavailable.

Enzymes that are not spatially represented in the model are treated as well-mixed and their rate-limiting effects are treated in a simplified manner. For these, we turned to transcriptomics and iBAQ proteomics data (30) and traditional FBA analysis using cobrapy (77) to predict whether they carry flux at the two time points considered. All essential reactions—those whose absence results in zero biomass accumulation or maintenance flux (i.e., cell death) in the GSMM of (43)—were assumed to carry flux. All others were permitted to carry flux only if we estimated their catalyzing enzyme(s) to be upregulated. To perform this estimation, we considered the median-centered (log2-scaled) iBAQ values of each enzyme across time points (or log2 TPM if the enzyme was not captured in the proteomics) as a proxy for abundance. We consider an enzyme to be upregulated if its abundance proxy value is at least half its maximum across time points.

In the presence of light, we assume that RuBisCO efficiency in the carboxysome is limited to a conversion rate of 4.7 mmol/gDW/hr, following (43), and that this ultimately limits growth (i.e., we assume that the growth medium is nutrient-replete). In the absence of light, we allow consumption of stored glycogen at a rate of up to 0.3 mmol/gDW/hr, consistent with dynamic FBA predictions from (43). All other parameters in the metabolic model are as in (43).

#### Post translational modification rates

Following the overall research design focusing on the central carbon metabolism under light disturbance, the Prk_2_ and Gap_4_ enzymes from CBC are particularly important, as the activities of these enzymes are determined by the redox state of CP12. In the presence of light (i.e. reduced state) Prk_2_ is not sequestered by CP12(oxidized), this results in a higher fraction of active Prk_2_. Active Prk_2_ converts RuP into RuBP in the cytosol by using ATP produced from the light reactions of photosynthesis during the regeneration stage of the CBC. RuBP enters the carboxysome where RuBisCO catalyzes its reaction with CO_2_ to form 3-PGA in the carbon fixation stage of the CBC. In this part of the CBC, RuBisCO-catalyzed reaction that occurs in the carboxysome is the limiting reaction. During the reduction stage of CBC, which follows carbon fixation, Gap_4_ catalyzes the formation of G3P by consuming NADPH produced primarily through the light reactions of photosynthesis. During the nighttime (i.e. oxidized state), however, Prk_2_ and Gap_4_ are sequestered by oxidized CP12 and they form an enzymatically inactive protein megacomplex of Gap8CP124Prk4, the dark complex, in the cytosol. A homologue structural model of the MED4 dark complex is shown in **Figure 6**. The formation greatly decreased the availability of Prk_2_ to catalyze RuBP formation and of Gap_4_ to catalyze G3P formation. In these reductant-limited conditions, the Prk_2_-catalyzed reaction in the cytosol limits carbon fixation. Hence, the metabolic flux is shunted to the oxidative pentose phosphate pathway and glycolysis to generate reductant NADPH for sustaining cellular functions.

**Figure 6.**
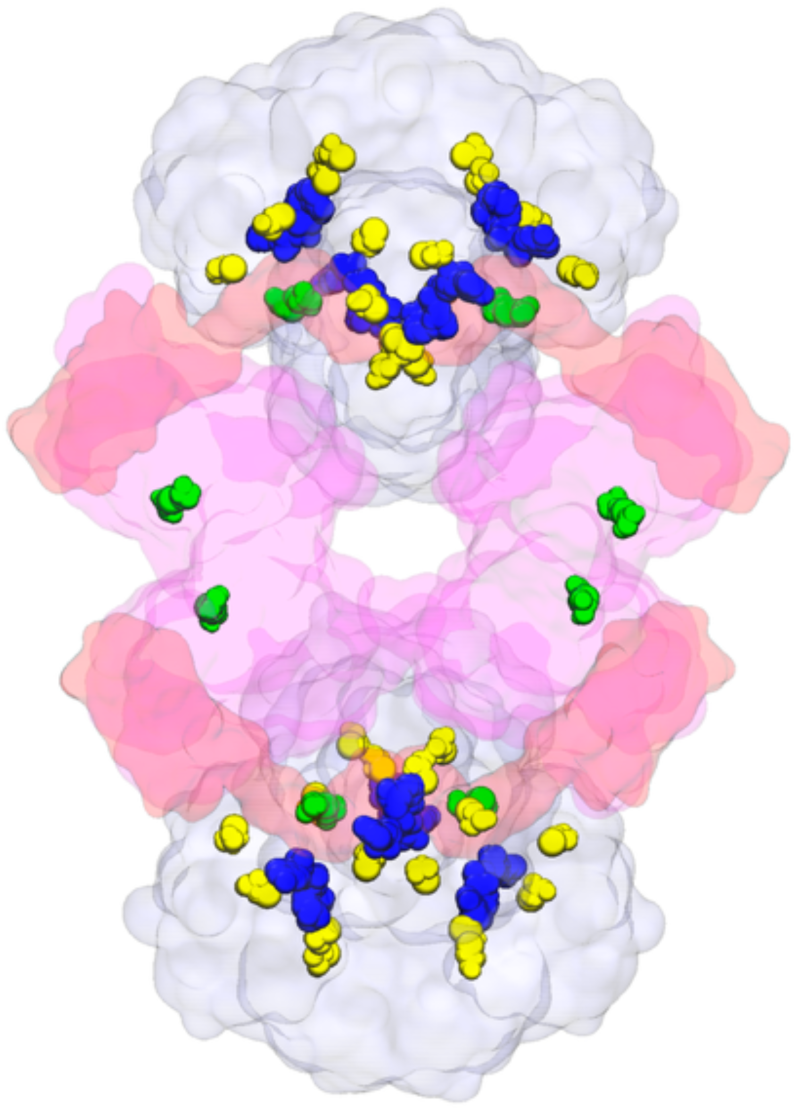
Structural representation of the MED4 dark complex with Gap8CP124Prk4 subunits: glyceraldehyde 3-phosphate dehydrogenase (a tetramer, Gap4) in grey, Phosphoribulokinase (a dimer, Prk2) in pink, and CP12 chains in red. Also highlighted are the pair of oxidized cysteines connected by a disulfide bond in green, cysteines in yellow, and nicotinamide adenine dinucleotide (NAD) in blue. This model was constructed as a homologue of PDB:6GVE which was experimentally determined from Thermosynechococcus Vestitus BP-1

Little is known about the rate at which light exposure induces oxidation changes in CP12 to disrupt the dark complex. The literature suggests that this process occurs over the course of several minutes in laboratory conditions (4, 78). This aligns with simple physiological considerations: assuming that the rate of oxidation state change is proportional to sinusoidally varying sunlight intensity, it is straightforward to show that a maximum CP12 oxidation rate (at local noon) of 0.005/s will oxidize 90-95% of CP12 within one hour after the beginning of sunrise. Under constant light, the same parameters yield a time of half-maximum CP12 oxidation of approximately two minutes. We have selected binding and dissociation rates for the dark complex such that these oxidative effects are the primary limiting factor, consistent with literature (4, 78).

#### Diffusion coefficients

Diffusion coefficients were computed for proteins and protein complexes (see **Tables 1** and **3**) using hydrodynamic estimates in HYDROPRO (58). Protein structures for these simulations were obtained using AlphaFold3. Diffusion coefficients for transcripts were estimated by assuming a hydrodynamic radius proportional to the cube root of transcript length, as in (79). Diffusion coefficients were initially estimated in water and rescaled. A correction factor was applied to account for reduced effective diffusion rates, which arise from macromolecular crowding interactions (80, 81) in prokaryotic cytoplasms (59). This method computes an effective diffusion rate that is decreased relative to free diffusion in proportion to the hydrodynamic radius (intuitively, larger molecules are more strongly affected by crowding).

### VI. Simulation, Analysis, and Data Visualization

We configured and deployed Lattice Microbes version 2.4 to enable 4D whole-cell simulations across a range of high-performance computing (HPC) centers. In addition, we configured and deployed VMD (82) with the Lattice Microbes and LAMMPS (56) plug-ins, enabling remote and local visualizations of large-scale whole-cell simulation trajectories.

We simulated four light conditions over two hours of simulation time with three replicates each. Our model captures the spatially localized RNAP-genome binding across the entire genome and resolved diffusion and complex formation for 15 key regulatory and enzymatic proteins using a spatial resolution of 20 nm. We capture enzymatically constrained metabolic fluxes across 994 reactions in a spatially coupled well-mixed model. We use a simulation time step of 8 μs and a write-out frequency of 1s. Visualizations of our simulations are available in the Supporting Materials. We also provide time series of aggregated mRNA counts, protein complex counts, and metabolic fluxes in Supporting Materials.

For each of four light conditions (Dark-to-Light, Light-to-Dark, Dark-to-Dark, and Light-to-Light), we simulated 100 replicates of a stochastic well-mixed chemical master equation (CME) model as a non-spatial baseline for genetic information processes. For the reaction diffusion master equations (RDME), for each of the four light conditions, we simulated 3 replicates of our whole-cell spatial model using independently generated DNA weave embeddings. Each simulation trajectory covered 120 minutes of cell time. Each RDME simulation leveraged NVIDIA A100 GPUs and CUDA with a typical wall clock runtime of approximately 80 hours for spatial simulations. Data plots were rendered using Matplotlib (83). We used pyLM (84) and ipython or jupyter notebooks (85) to assist with constructing and integrating the model components before running simulations on the HPC systems using SLURM (86). Finally, additional visualizations and analysis of the tomogram were performed with ChimeraX (49).

## Results

### The first 4-D whole-cell model of a photoautotroph reveals spatially resolved metabolic phenotypes in CBC regulation

The perturbation approach allows us to simplify the biology phenomena and follow the autotroph dynamics after light disturbance at selected light variations. Here, we present the two distinct metabolic phenotypes of an autotroph as illustrated in **Figure 1** and **Figure 2**, with a spatially resolved whole-cell model. We focused the spatial component on the primary energy metabolism of MED4, with special emphasis on the carbon Lixation regulated by redox changes in light conditions. Dynamically, we simulated the expression for a subset of 15 genes within the MED4 genome (**Table 1**). These genes correspond to cytoplasmic enzymes from key metabolic pathways in central metabolism, transcription factors which modulate nutrient uptake, or have shown notable changes expression proLile under varying lighting conditions.

After we set the coefficients, variables, mathematical expressions, and located the intracellular reactions, we presented the first whole-cell model (WCM) of a photosynthetic autotroph using Lattice Microbes (LM) software of a laboratory-grown *Prochlorococcus marinus* MED4. Spatially, LM discretizes the cell volume on a cubic lattice to represent boundaries for large subcellular structures and compartments. As shown in **Figure 6c**, the WCM of MED4 that build on a high-resolution (10 nm) cryo-electron tomography includes key spatially resolved components of thin-layered thylakoids under the outer membranes and the two spherical carboxysomes. We next brought the “molecular biology of the central dogma” into the LM in that the transcription and translations are spatially resolved in the WCM. LM expands the genome-scale metabolic modeling with cell-scale dynamics of key expressed proteins and mRNA transcripts, as well as RNAP binding to a polymeric model of whole genome.

Figure 7 depicts the two initial conditions of the model corresponding to dawn (6AM; Figure 8A) and dusk (6PM; Figure 8B). The former represents a phenotype with a highly oxidized pre-sunrise intracellular environment that promotes the formation of an inactive Gap_8_CP12_4_Prk_4_, the dark complex (magenta spheres), in the CBC. The latter represents a phenotype with a highly reduced pre-sunset intracellular environment in which Prk_2_ (green spheres) and Gap_4_ (navy spheres) are active to catalyze metabolism in the cytosol.

**Figure 7.**
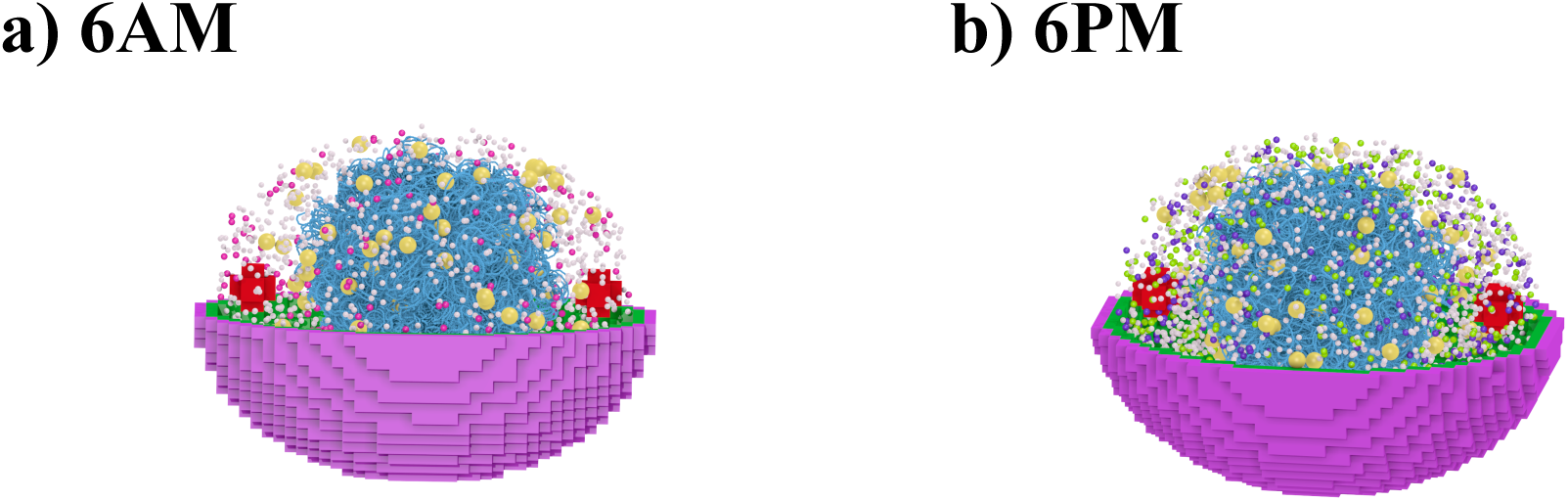
Spatial representation of key enzymes in the CBC in a whole-autotroph model at a) 6AM at dawn and b) 6PM at dusk. Prk2 (green), and Gap4 (navy), dark complex (magenta), ribosome (yellow), thylakoid (green), cell wall (purple), carboxysomes (red), and weaved DNA (blue).

**Figure 8.**
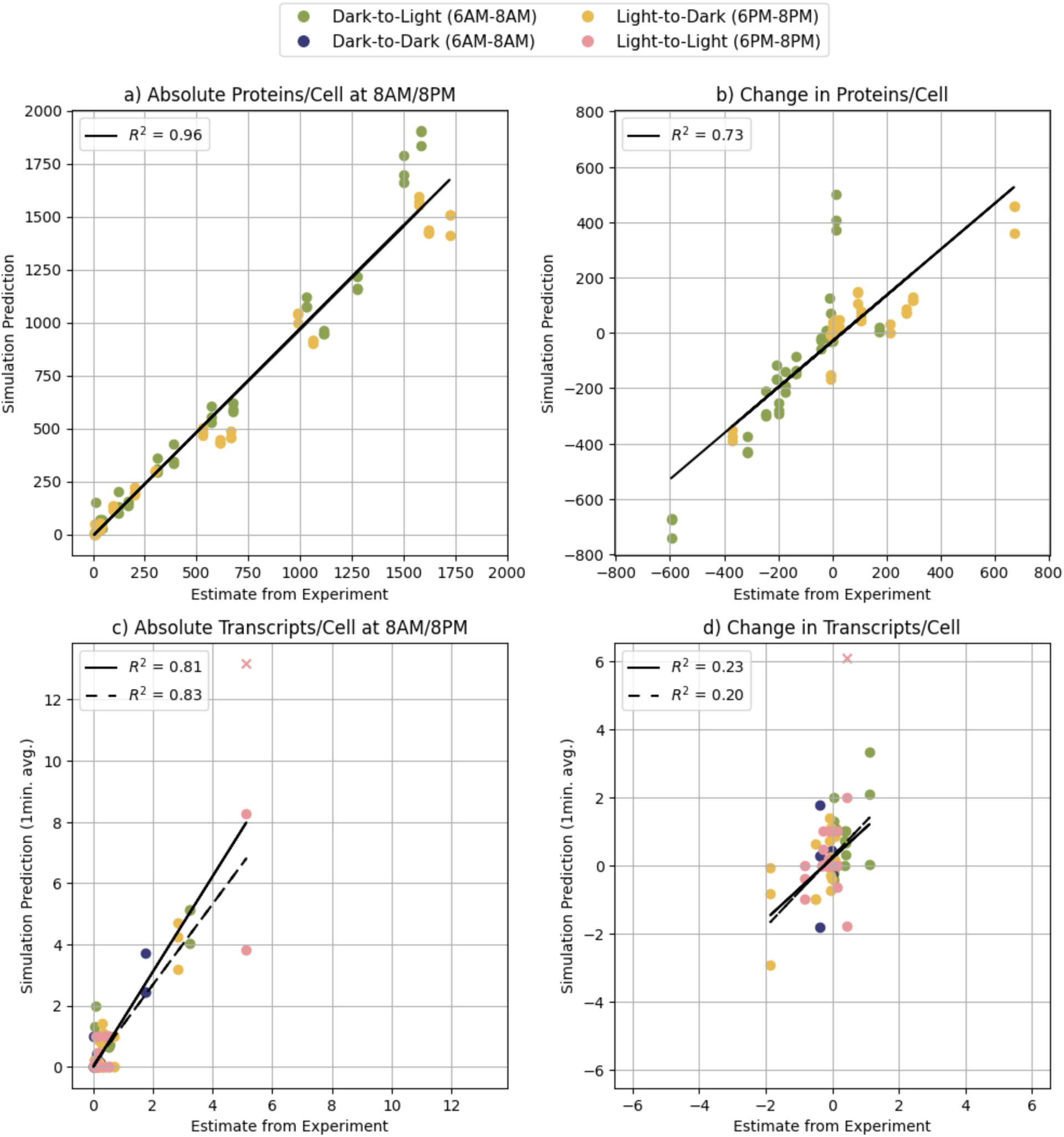
Comparison of experimentally estimated and protein and transcript counts simulated by LM WCM over 2 hours with 8 microsecond time steps. a) Comparison of absolute counts at end of simulation period (8AM for Dark-to-Light scenario and 8PM for Light-to-Dark scenario). b) Comparison of change in protein counts over the two-hour simulation period. Simulated protein counts include all proteoforms and complexes. c-d) Comparisons for absolute transcript count and change in transcript count. Simulated transcript counts are computed as the average number of transcripts observed over a period of one minute to account for the high stochasticity of gene transcription. For transcript counts, genes with no captured transcripts are omitted. An outlier in simulated transcript counts is observed in the case of prk for the Light-to-Light scenario (indicated by a cross symbol). We performed linear regression with (solid lines) and without (dashed lines) this point and found comparable R^2^ values. Proteomics data is provided for the Light-to-Light and Dark-to-Dark scenarios.

### Realistic predictions of the simulated mRNA and protein counts representing different phenotypic metabolic states under light variations show good agreement with experimental data

The Lattice Microbes for MED4 model integrates transcriptomics, proteomics, and cryo-electron tomography with Brownian dynamics simulations for DNA packing, Markov models for rate parameter estimation, and structure-based diffusion simulations. This culminates in a 20 nm resolution, 8 μs time-step whole-cell model (WCM) simulation of spatial genetic information processing and carbon fixation regulation within the cytoplasm of the world’s smallest autotroph. A represented movie is provided in the Supplement.

We executive the WCM simulations under four light conditions following the overall research design described in Figure 2. We checked the implementation performance of the model by comparing the protein counts and transcript counts for the 15 genes listed in **Table 1** from the simulations against the single cell count estimates from our multiomics experiments (as described in the previous section) in Figure 8. These experiment measurements include bulk proteomics for the Dark-to-Light (dawn) and Light-to-Dark (dusk) scenarios (Figure 8a**,b**) whereas bulk transcriptomics for all four scenarios include two light perturbations (Figure 8c**,d**,). We started with the initial counts for both proteins and transcripts at 6AM (dawn) or 6PM (dusk) and then compared the simulated and experimentally measured protein and transcript final counts after two hours at 8AM or 8PM. We find good agreement both in estimated absolute counts (R^2^ > 0.8;) for proteins (Figure 8a) at 8AM (2 hours after dawn) or 8PM (two hours after dusk). We also find good agreement with the estimated change in protein counts over the two hours between the simulated and the experimentally measured ones (R^2^ = 0.73; Figure 8b).

For transcript counts, we compared the initial and final counts at dawn (Dark-to-Light) and at dusk (Light-to-Dark) **in** Figure 8c**, d**, as well as two light perturbation conditions (Dark-to-Dark) and (Light-to-Light). In the case of simulated transcript counts and the changes over two hours, we compute a one-minute average to better account for the extreme stochasticity of transcription. We omitted from transcriptomics comparison any gene for which the experimentally estimated initial and final transcript count was zero. We find good agreement with the absolute counts (Figure 8c). The change in transcript counts, however, is less satisfactory (R^2^ = 0.2, Figure 8d) due to the need to round the number of initial transcript counts.

Specifically, we depicted the simulated initial and final transcripts counts for the 15 spatially tracked gene transcripts and proteins (aggregating monomers across tracked complexes) for the Dark-to-Light (dawn) and Light-to-Dark (dusk) scenarios over two hours in Figure 9, for which we have collected both proteomics and transcriptomics data. In Figure 9a, the *prk* gene has the highest initial and final transcript counts under both conditions. In contrast, and despite their low transcript counts, glycolytic or redox enzymes such as gene products for *gap2*, *eno*, *glpX*, and *petH* have the most protein counts over 1000 in Figure 9b. For the two perturbed Dark-to-Dark and Light-to-Light, we provided them in **Figure S4 a, b**, respectively. These visuals signify that the WCM model captured distinctive phenotypes in response to light variations from the same genome in an autotroph

**Figure 9.**
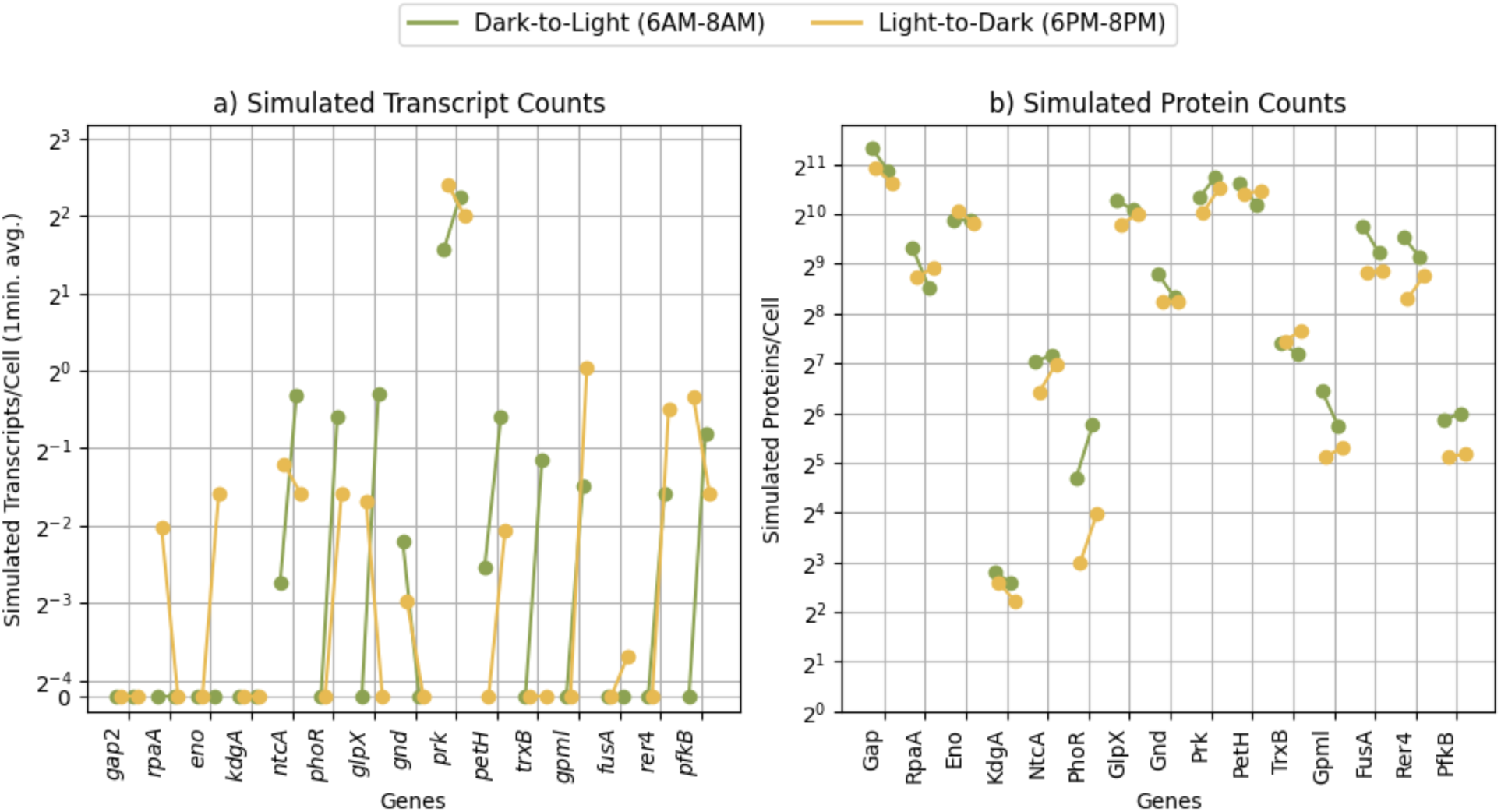
Comparison of initial and final states. In terms of transcript counts in a) and protein counts in b) for selected 15 genes and gene products in the scenarios at dawn (Dark-to-Light) or at dusk (Light-to-Dark) by LM WCM over 2 hours with 8 microsecond time steps. a) Initial and final transcript counts; simulated transcript counts are computed as the one-minute time average of transcript counts. b) Initial and final protein counts. Note that protein counts include the proteoforms and complexes for the dark complex (Table 3). In both panels, the left endpoint of each line segment indicates the initial count and the right endpoint indicates the final count after two hours of simulation.

In summary, we observed large variation in transcript counts across stochastic replicates. This is likely due to the extremely low transcript count per cell---often less than a single transcript per cell, on average, in line with previously published measurements in bacteria (87). This variation at the mRNA level is largely absent at the level of proteins in our simulation. Simulated protein counts per cell are on the order of hundreds or thousands, in good agreement with our proteomics-based measurements and in line with direct measurements in *E. coli* (MED4 is approximately half the volume of *E. coli*) (87).

### The temporal profiles of CBC enzyme counts from the CME and RDME models are more robust than those of transcript counts, due to post-translational regulation

We compared the outcomes from CME and RDME by analyzing the output trajectories. The largest difference we observed between the models was in the transcript counts for *prk* (**Figure S2**). Because the RDME incorporates additional stochasticity by explicitly modeling the diffusion of individual RNAP molecules to lone transcription sites, there is substantially more temporal and replicate-to-replicate variation in transcript counts as compared to the CME model. Moreover, the RDME naturally incorporates the possibility for RNAP to rebind the promoter region of gene following transcription, which occurs at a rate higher than would be inferred under a well-mixed approximation. This has the largest effect on highly expressed genes (*prk* is the most highly expressed gene in our study). The average transcript count is up to 50% higher in the RDME than the CME simulation, despite both models using equivalent rate parameters.

At a protein level, the temporal profiles of protein counts from the CME and RDME models are much more similar (Figure 10), indicating that the transcriptional and translational noise is buffered by post-translational effects. Interactions between proteins give rise to rich dynamics in response to changing light and redox states, including non-monotonicity in the counts of protein complexes. This arises because of the interplay between redox-driven complex assembly and disassembly with protein synthesis and degradation. The sequestration and release of the active enzymes in response to changing light conditions occurs on similar time scales approximately within an hour in both models, consistent with previously reported observations on the timescale of carbon metabolism activation and inactivation (4, 78).

**Figure 10.**
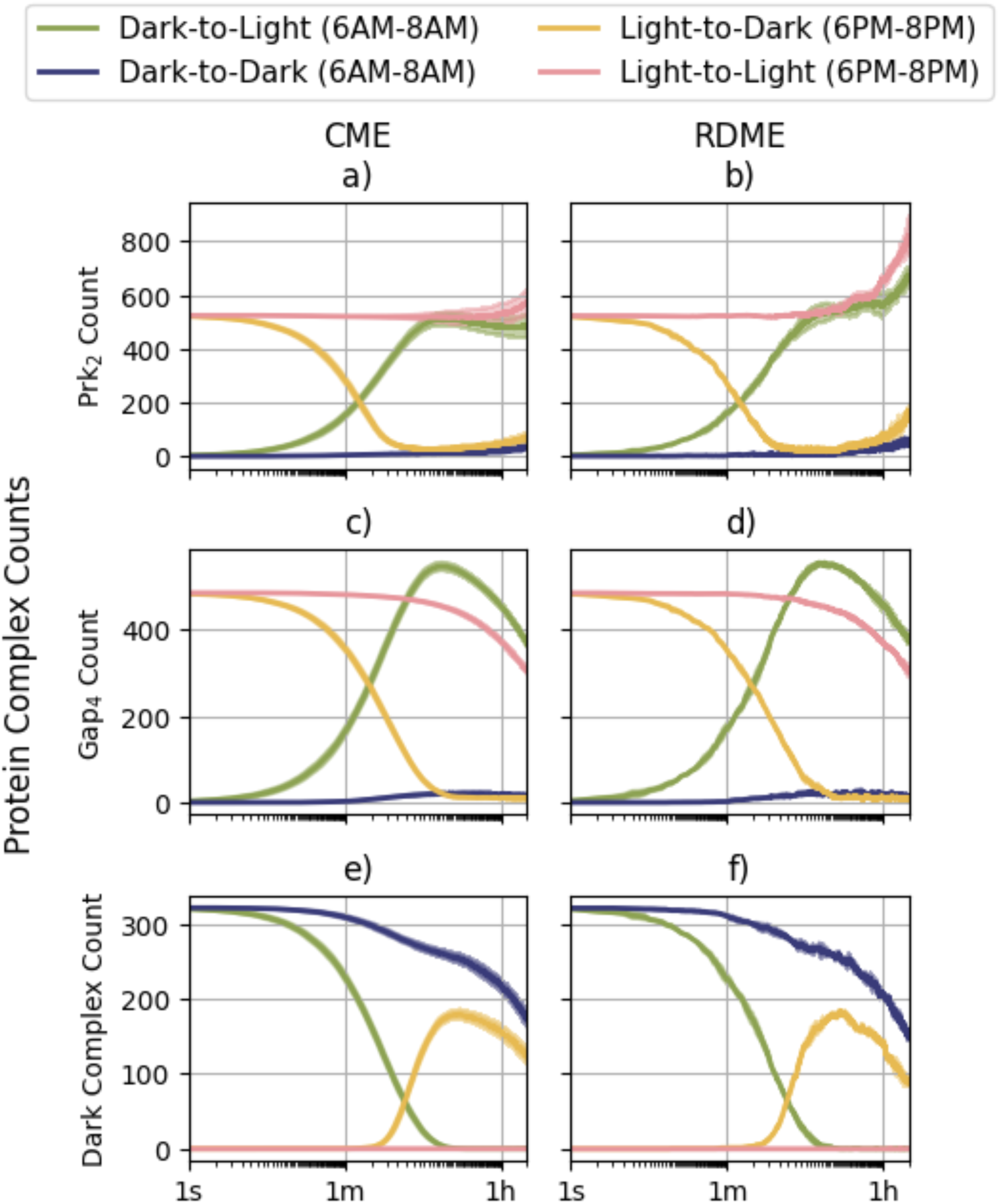
The impact of spatial heterogeneity on formation of the dark complex. a,c,e) Simulation outputs from a well-mixed CME model of metabolic gene transcription (100 replicates per light condition). b,d,f) Simulation outputs from the spatially resolved whole-cell model (3 replicates per light condition). Each row depicts a key protein complex: a,b) Prk2 dimer; c,d) Gap4 tetramer; e,f) Dark complex, Gap8CP124Prk4. Colors indicate the simulated light condition. Average protein count key complexes are represented by solid lines. Shaded regions correspond to one standard deviation of replicate values. The same binding parameters, up to a diffusion correction, are used for the simulations in both panels.

The phenotypic metabolic states in CBC are manifested by the redox proteoforms and molecular assembles such as Prk_2_, Gap_4_, and the dark complex, Gap_8_CP12_4_Prk_4._ High abundance of active Prk_2_ enzyme (**Figure 10a, b** for CME and RDME, respectively) corresponds to the light (reduced) conditions, while being low abundance in the dark (oxidized) conditions. The Gap_4_ enzyme initially increases in a similar manner as the initial population of dark complexes dissociates (Figure 10 **c, d** for CME and RDME, respectively). At later timepoints, excessive Gap is degraded and not replaced (the *gap2* gene is only minimally expressed as discussed in subsequent sections). In the absence of light, the enzymatic activities of Prk_2_ and Gap_4_ in the dark complexes are sequestered by oxidized CP12, and the protein counts of the dark complex grow non-monotically under light-to-dark condition as the constituent enzymes are sequestered on short timescales and degraded on longer timescales (Figure 10 **e, f** for CME and RDME, respectively).

While both the CME and RDME simulations predict a small stoichiometric imbalance between Gap_4_ and Prk_2_ close to an hour, this imbalance is amplified in the RDME simulations. This excess Prk_2_ cannot be fully sequestered because there is insufficient Gap protein; this leads to metabolic inefficiencies that we explore in subsequent sections.

### Redistribution of metabolic flux for survival in response to light variation

When exposed to light, the light reactions of photosynthesis generate ATP using the proton gradient established across the thylakoid, as captured in our simulations (**Figure 11a**). In the absence of light, ATP generation primarily takes place in the periplasm (**Figure 11b**), and energetic requirements are ultimately derived through glycolysis and the oxidative pentose phosphate pathway (PPP) from glycogen reserves accumulated during previous light exposure (**Figure 11c**). The PPP enzyme GndA produces Ru5P, which can be phosphorylated by Prk_2_ as part of the CBC (**Figure 11d**). Tkt catalyzes a reversible reaction which produces either G3P or ribose-5-phosphate (R5P), a precursor for Ru5P, in opposite reaction directions. When light is abundant, Tkt has a higher propensity to run in the direction which favors production of R5P; in dark conditions this is reversed, favoring the production of G3P (**Figure 11e**). In light, carbon fixation is essential for biomass growth; in darkness, carbon fixation represents an inefficient usage of cellular resources, inhibiting biomass growth (**Figure 11f**). This progression is summarized schematically in **Figure 11g**.

**Figure 11.**
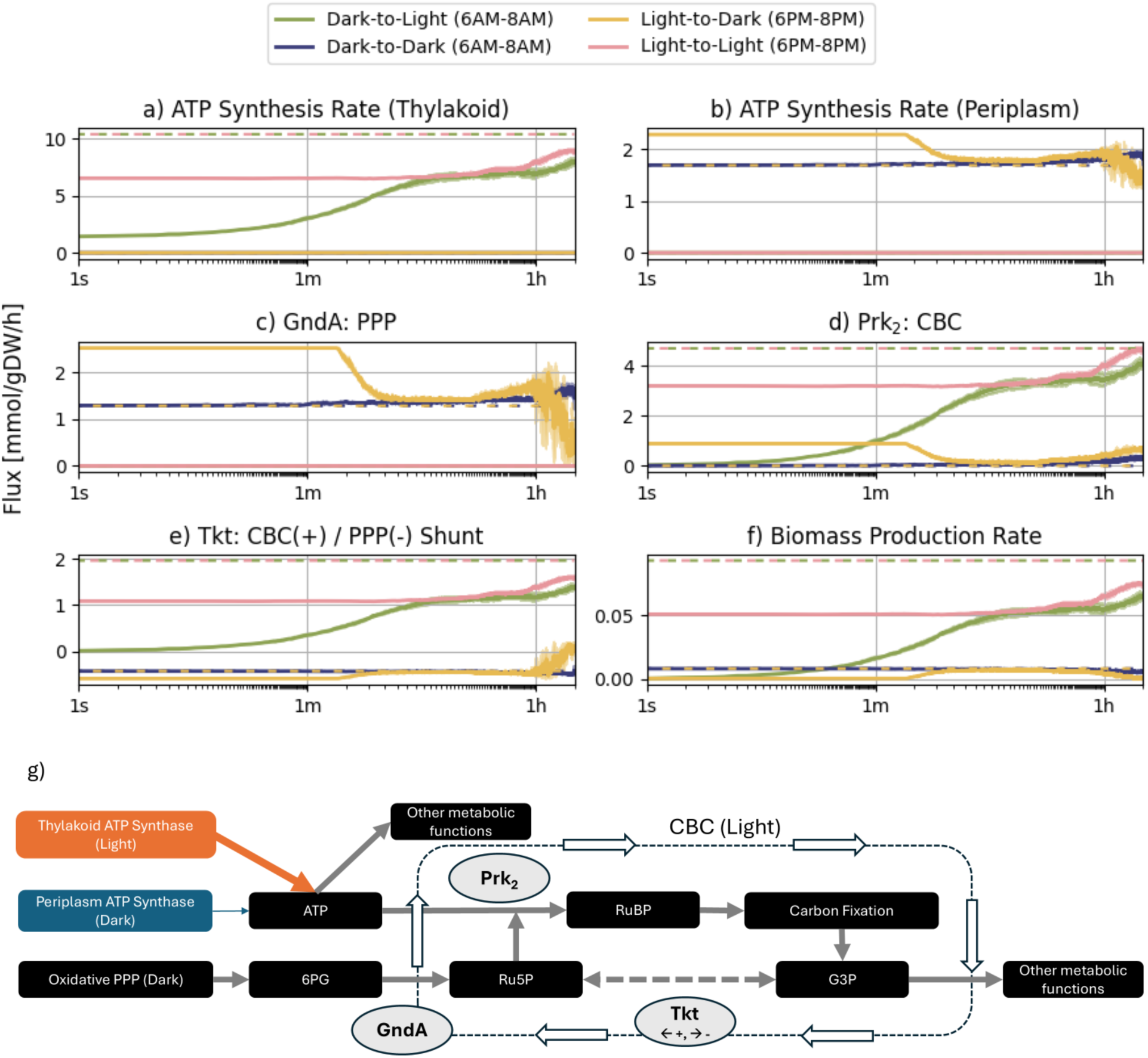
Metabolic flux time series representing critical metabolic processes. (a-f) Colors indicate light conditions. Solid lines represent average flux values across three replicates with shaded regions representing the standard deviation of replicate values. Dashed lines indicate baseline fluxes obtained from unconstrained flux balance analysis. Each panel indicates a different metabolic process: a) photosynthetic ATP synthesis in the thylakoid, b) glycogenolysis-driven ATP synthesis in the periplasm, c) pentose phosphate pathway (PPP) activity as represented by the GndA catalysis of Ru5P formation, d) carbon fixation in the Calvin-Benson cycle (CBC) n represented by the phosphorylation of Ru5P by Prk2 e) CBC vs PPP activity as captured by the reversible production conversion of G3P to Ru5P by transketolase, f) biomass formation (see Figure S1). Panel g depicts a simplified schematic illustrating how the reactions in panels a-e are interconnected (see text for further details).

We compared the reaction fluxes in our RDME simulations to optimal flux distributions obtained by flux balance analysis in the absence of RDME-derived enzyme constraints (**dashed lines in** Figure 11). In our RDME simulations, we observed that incomplete inhibition of Prk_2_ early in the Light-to-Dark transition leads to increased demand for ATP (to phosphorylate Ru5P) relative to the optimal generation rate. We also observed reduced biomass accumulation rate. After approximately two minutes, enough Prk_2_ is sequestered in dark complex (or its intermediates) that it becomes rate limiting for the phosphorylation of Ru5P, and carbon fixation is inhibited. At this point, ATP demands are decreased and biomass accumulation increases, both achieving values near the unconstrained, optimal fluxes. On longer timescales (beyond one hour), fluctuations in enzyme availability and stoichiometric imbalance in dark complex constituents (Figure 10) due to stochastic effects lead to transient decreases in metabolic efficiency for the Light-to-Dark scenario.

The biomass accumulation is reduced in the no-light conditions, particularly in the Light-to-Dark scenario in which carbon fixation and the PPP are far from their optimal values due to excess Prk_2_. Despite very low biomass accumulation rates, we observed that at all time points in our simulation, and in all replicates, the metabolic maintenance reaction maintains its maximum flux, indicating continued cell survival. Cell death and biomass depletion in darkness are expected at low rates in MED4, though this depends on cell-to-cell variation in glycogen stores. In our simulations, we have opted to consider cells with sufficient glycogen stores to avoid cell death during the simulation period (**Figure S2**). The biomass accumulation rate is consistent with values reported previously in the literature (43). Moreover, daytime biomass accumulation rates of between 0.05 and 0.1 g/gDW (as seen in the simulation results of **Figure 11f**) are consistent with a doubling time 24 hours, given that significant biomass accumulation occurs only during the daytime (16, 43) and assuming exponential growth.

### Emergence of subcellular structures and local molecular diffusions sustain metabolic phenotypes under light disturbance

*Light perturbations (as defined in* Figure 2*) revealed a tightly coupled response between the thylakoid light reactions, cytosolic redox state, and carboxysomal carbon fixation*. In the Dark-to-Light case, photosynthetic flux was initially limited not by light availability but by the cell’s capacity to process downstream metabolites, because Prk_2_ and Gap_4_ were largely sequestered in the redox-regulated dark complex. As light-induced redox changes rapidly drove dark-complex disassembly in the cytosol, the concentration of active Prk_2_ and Gap_4_ increased, relieving this bottleneck. Consequently, carbon fixation in the carboxysomes rose from its initially suppressed state and aligned with the enhanced photosynthetic throughput, indicating that the limiting step shifted from enzyme availability to substrate supply as the system transitioned to the illuminated state (Figure 12).

**Figure 12.**
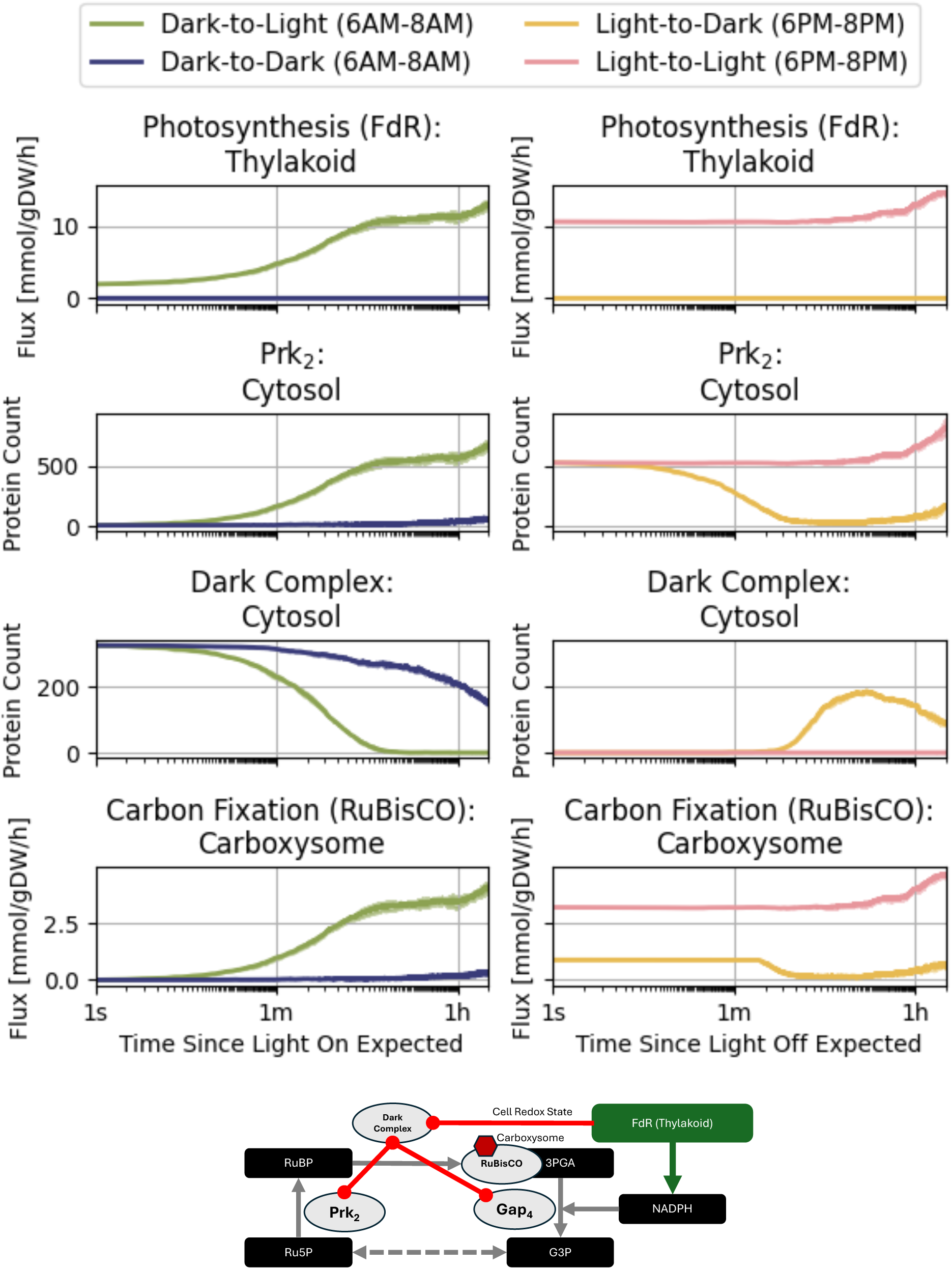
Coupling between the thylakoid and carboxysomes via redox-modulated reaction diffusion processes in the cytosol. Two circadian-aligned simulation scenarios are considered: Dark-to-Light (green, left column) and Light-to-Dark (yellow, right column), along with their perturbed counterpart scenarios Dark-to-Dark (blue, left column) and Light-to-Light (pink, right column). In each plot, the initial states of both the circadian-aligned and circadian-perturbed time series coincide with the circadian-perturbed initial condition. Discontinuities in the flux (the time derivative of the metabolite production) arise because the circadian light changes are modeled as instantaneous. In the schematic at bottom, the redox coupling between FdR in the thylakoid and RuBisCO in the carboxysome, mediated by cytosolic enzymes and complex formation, is depicted in simplified form (see text for further details).

The reverse transition, from Light-to-Dark, displayed a different hierarchy of control. Immediately upon light removal, thylakoid light reactions ceased, causing an abrupt drop in photosynthetic flux and a concomitant decline in carbon fixation. Nonetheless, residual activity of CBC enzymes allowed carbon fixation to persist at a level that remained sufficient to inhibit complete shutdown of growth and maintenance processes. Further suppression of carbon fixation required an additional regulatory step: *sequestration of Prk_2_ and Gap_4_ into the dark complex as the cytosolic redox potential became more oxidizing in response to the halted light reactions*. Only when the abundance of active Prk_2_ fell below a critical threshold, approximately 100 s after light removal, did carbon fixation decline further toward its dark steady state (Figure 12). These simulations show that redox-modulated complex formation in the cytosol acts as <u>a delayed, enzyme-level brake</u> on carbon fixation, complementing the immediate, light-dependent control exerted by the thylakoid and thereby coordinating energy capture and carbon assimilation across compartments in a cell over multiple timescales.

In the Dark-to-Dark scenario, there is no flux through the FdR reaction at any time point (due to the lack of light). Carbon fixation remains low, but the dark complex and its constituents degrade over time. During this degradation, some small amount of transient carbon fixation occurs (on hour-timescales). Similarly, photosynthetic light reaction and carbon fixation reaction fluxes in the Light-to-Light scenario remain stable for almost one hour, but stochastic effects led to increase production of Prk_2_ and a subsequent increase in both fluxes (Figure 12).

## Discussion

### I. Tracking the outcomes from environmental perturbations with synergistic efforts of experimentation and modeling enabled the first 4D whole-cell simulations of an autotroph

MED4 is particularly well suited as a target for the first whole-cell–style model of an autotroph (i.e. microbial organisms that produce their own food from inorganic materials and sunlight, for example). Its small size, less than one micron in diameter, enables cryo-ET imaging (17) of the entire cell without destructive milling. Its tomogram yields a relatively small simulation domain. Despite the merit of reducing the computational cost, MED4 is genetically intractable, limiting direct perturbation experiments and impeding classical approaches to studying its regulatory pathways (88). We therefore relied on *environmental perturbations* such as illumination variations with a multimodal, whole-cell–oriented strategy to achieve a comprehensive modeling of its metabolisms that combines transcriptomics, proteomics, structural modeling, genome annotation, and cryoET-based morphology, supplemented by homology-based inference from well-studied relatives such as *Synechococcus elongatus*. This approach is applicable not only to MED4 but also to other non-model organisms of potential biotechnological or biothreat relevance, where genetic tools may be unavailable.

The perturbation-based approach with synergy of experimentations generated multimodal datasets from the same culture samples in a controlled setting. The datasets generated in this work capture the causal relations across omics layers. It enabled a comprehensive modeling of events across disparate time scales connecting light-induced redox changes in the metabolic states, transcriptional responses, dark-complex formation, and CBC fluxes. We deliberately selected a minimal gene set informed by -omics experiments and enzyme parameters to test a specific whole-cell–scale hypothesis that to understand complex cellular responses under light disturbance. Fast time light-dependent redox PTMs that regulate the structural assembly of the dark complex, control the metabolic flux at a conserved regulatory node of the CBC in cyanobacteria. A key feature of this model is its incorporation of redox PTMed dark complex, a critical process in photosynthetic regulation (89). A key feature of this model is its incorporation of redox PTMed dark complex, a critical process in photosynthetic regulation (89). We demonstrate consistency between our model and existing experimental and modeling literature, including in the biomass accumulation rates and key light/dark transitions in the metabolic state of the cell (4, 16, 43) (see Figures 10**, 11**). Our model provides mechanistic insight that the redox-modulated assembly of the dark complex transmits environmental signals from light-dependent reactions at the thylakoid to light-independent central carbon metabolism at the carboxysome, triggering carbon fixation when light is available or the oxidative pentose phosphate pathway to generate reductants at light-limited circumstances (Figure 12). Accompanying this manuscript, we have released reusable datasets in parameterizing this model to aid and evaluate future whole-cell modeling efforts (90).

### II. Organelles solve the need for spatial organization of biological functions under the light-energy limitation, but require intracellular coordination

*Prochlorococcus marinus* MED4 presents a particularly stringent test case for spatially resolved modeling because of its extremely small size and correspondingly fast diffusion, which places nearly all intracellular chemistry in a reaction-controlled regime. Organelles and microcompartments solve a fundamental need for spatial organization of biological function despite the apparent homogeneity of such a small cell. Thylakoids are required to generate proton gradients and reductants such as NADPH, whereas carboxysomes are required to concentrate CO₂ and RuBisCO, preventing oxygen from competitively inhibiting carbon fixation. The spatial compartmentalization which establishes these chemical gradients is independent of the day–night cycle and helps explain why MED4 retains elaborate internal organization even under constant light conditions. This segregation of metabolic function raises a new challenge for the cell, however: information must be passed between compartments to harmonize cellular function.

Our model elucidates redox PTMs in the cytosol as a mechanism transmitting the reduced photosynthetic activity in the thylakoid to inhibiting carbon fixation in the carboxysome.

We show that under light perturbations when photon and electron influx in the thylakoid change rapidly, key enzymes in the CBC are sequestered and differentially regulated by redox-dependent post-translational modifications (redox PTM), providing a mechanism to couple environmental light information to the regulation of carbon fixation with molecular mechanisms (91). In the absence of light, the CBC enzymes glyceraldehyde-3-phosphate dehydrogenase (Gap_4_) and phosphoribulokinase (Prk_2_) are inhibited by the redox active CP12 protein that sequesters and binds these enzymes into the Gap_8_CP12_2_Prk_4_ *dark complex* within minutes (92). Sequentially, two oxidized CP12 protein copies bind to the Gap_4_ tetramer before two Prk_2_ dimers bind to CP12 to bridge two Gap_4_CP12_2_ complexes, forming the full dark complex (Figure 7). In contrast, the dark complex quickly dissociates under light conditions, releasing active P4k_2_ and Gap_4_ enzymes and turning on complex enzyme regulation (21).

Dark complex formation or disassociation depending on its redox state takes place on the scale of minutes on demand, while other genetic processes take place on hour-scales leading to a more robust, sustained change in metabolic state. Thus, the spatiotemporal coupling of reactions and PTMs across multiple time scales and between spatially separated compartments shapes the metabolic phenotypes of the cell. Understanding the dynamic behavior of these multienzyme megastructures can provide insights into metabolic regulation and could guide bioengineering interventions for pathway efficiency (93).

The separations of organelles allow sequestered biochemical reactions taking place in varying time scales. For example, fast reactions in photosynthesis are sequestered in a thin layer of thylakoids close to the outer membrane. Part of the slowest reactions in CBC for carbon fixation are sequestered in the carboxysomes for accessing highly concentrated CO_2_. By including the molecular insights from PTM-guided assembly of protein metacomplexes that act as “redox switches”, we bridged the knowledge gap of connecting light reactions in photosynthesis and light independent CBC carbon fixation with redox PTM-mediated complexation of key regulators and enzymes, and their spatially resolved impact on genome-scale metabolism.

### III. PTMs drive robust responses to environmental changes in the first whole-cell model of a photoautotroph

In this study, we observed large stochasticity in prk and gap transcript counts, but a robust response of protein concentrations of the metabolically active Prk_2_ dimer and Gap_4_ tetramer whose monomers they encode. The catalytic activity of these enzymes is severely inhibited by the light-dependent redox modification of the quickly diffusing CP12 protein by forming a dark complex. Well-mixed CME simulations and spatially heterogeneous RDME simulations showed marked differences in mRNA dynamics, but qualitatively similar proteoform counts over time. This result demonstrates how post-translational regulation of enzymes through protein-protein interactions, redox modification, and degradation overcome transcriptional stochasticity induced by spatial heterogeneity. That is, molecular megacomplex formation, such as dark complex, serves as a mechanism to maintain robust regulation of cyanobacterial energy metabolism in a heterogeneous intracellular environment over a diel cycle, bridging a critical knowledge gap about the emergence of diverge phenotypes from multiple metabolic pathways under redox stress in genotype to phenotype research.

Our simulations highlight a form of spatiotemporal canalization that extends classical concepts of buffering environmental stimuli into the development of traits (94) or phenotypes in an organism. There are large fluctuations in transcript count, but this stochastically varying transcriptional state is canalized in the phenotypic landscape toward a robust metabolic state through post-translational modification, protein-protein interaction, and protein degradation. By comparing the outputs of our 4D spatiotemporal kinetic model to an equivalently parameterized stochastic model, we see that the noisy diffusion of RNAP to an individual promoter region amplifies transcriptional noise, but that this noise is buffered at the level of active enzyme abundance, PTM-induced interactions, and of the metabolic fluxes these enzymes constrain. This integration of spatial stochastic kinetics into metabolic modeling is therefore essential for a complete understanding of how energy-limited, spatially organized cells maintain robust function under light variation.

Our model thus serves as a baseline for exploring further perturbations. For instance, MED4 has been used to study cyanophage-cyanobacteria interactions (17, 95), which are important for global nutrient cycling (96) and have potential bioengineering applications (97). Of particular interest is the incorporation of viral auxiliary metabolic genes such as CP12 (98) and the impact of circadian regulation or infection on cell morphology and metabolism. More broadly, this work establishes a data-driven, spatially explicit simulation methodology that advances fundamental understanding of how environmental sensing, genetic information processing, morphology, and post-translational regulation are integrated within photosynthetic cells.

### IV. Organism-agnostic requirements for whole-cell spatiotemporal modeling of responses to perturbation

Whole-cell models (WCMs) (99–101) integrate heterogeneous data and multiscale processes to reveal how cellular phenotypes emerge from stochastic molecular interactions. Our work extends this paradigm into a regime that has been largely inaccessible to existing WCMs: an energy-limited, photosynthetic autotroph experiencing light perturbations, with explicit spatial structure and redox-regulated protein megacomplexes. While most prior whole-cell and virtual cell models (100, 101) focus on heterotrophs and rely on well-mixed kinetics, our framework couples RDME-based reaction–diffusion dynamics. Because mRNA copy numbers are extremely low (often of order one or less per gene per cell) it is appropriate to model transcription and translation as stochastic processes whose reaction–diffusion dynamics directly shape information-processing rates (54, 87). Our cell-scale spatial model of a minimal photoautotroph shows how a small number of regulatory proteins such as CP12 can modulate genome-scale metabolism, and how transcriptional noise is buffered by protein degradation, complex formation, and redox modification. Together, these features demonstrate how WCM principles can be adapted to build spatially explicit, light-regulated “digital twins” of photosynthetic cells and provide a general framework for annotating protein function in the context of specific cell states.

In this work, we incorporated light-driven redox regulation, post-translational modifications, and constraint-based metabolic fluxes in a unified, spatially resolved WCM simulation. Incorporating a key representation to include redox-regulated PTM network with the assembly of the dark complexes into a whole-cell–style model is a key departure from earlier Lattice Microbes and WCM efforts, which generally treat enzymes as well-mixed species and rarely resolve photosynthetic regulatory megacomplexes.

In doing so, it provides a mechanistic bridge from genotype to phenotype by representing not only gene expression, but also proteoforms, multienzyme assemblies, and their regulation under different environmental conditions. This cross-scale coordination, central to WCMs, allows us to simulate cellular processes, perturb parameters, and predict regulated behaviors while explicitly visualizing how molecules diffuse and interact within a crowded, compartmentalized interior (59, 102, 103).

## IV. Conclusion and Future work

In this work, we have designed and executed concurrent perturbation experiments to generate multimodal datasets from the same sample cultures, which we have applied to generate the first whole-cell model of photoautotroph. We developed our model to probe the regulatory processes that enable communication between metabolically important subcellular compartments. We showed that biochemical regulatory interactions between these spatially separated compartments span three orders of temporal magnitude within a micron-sized organism, providing mechanistic insights for temporally resolved phenotyping.

The high-quality, curated datasets from this work and our model can serve as a useful tool for computer-aided augmentation and engineering of cells for bioeconomic synthesis (13, 22) and new discoveries, allowing researchers to probe cellular dynamics that are otherwise challenging to study experimentally. Our approach provides an extensible framework for modeling under-studied, genetically intractable organisms. This is a critical capability for addressing emerging biothreats and harnessing novel biochemical properties in non-model organisms.

## Data Availability

Raw proteomics measurement data files are openly accessible for download at the Mass Spectrometry Interactive Virtual Environment (MassIVE) community repository under the following data accession MSV000100371 (68). Primary RNA-seq data are openly accessible for download at the Gene Expression Omnibus (GEO) community repository under the data accession GSE314951 (73). Processed results files for proteome and transcriptome analyses, and LM simulation videos, are openly accessible for download at the PNNL NW-BRaVE Zenodo community (30) under the DOI: 10.5281/zenodo.18022273. Download includes experimental metadata, normalized quantification data, computed outputs, experimental metadata, and annotated results.

## Software Availability

The software and model schema described here is openly available from the NW-BRaVE GitHub repository, (https://github.com/NWBRaVE/Lattice_Microbe_Prochlorococcus), under “Lattice Microbe Prochlorococcus” supporting process transparency and can be formally cited under 10.5281/zenodo.18025765 (90) in alignment with source code community best practices.

## Acknowledgments

The research described in this paper is supported by the NW-BRaVE for Biopreparedness project funded by the U. S. Department of Energy (DOE), Office of Science, Office of Biological and Environmental Research, under FWP 81832. A portion of this research was performed on a project award (Enhancing biopreparedness through a model system to understand the molecular mechanisms that lead to pathogenesis and disease transmission) from the Environmental Molecular Sciences Laboratory, a DOE Office of Science User Facility sponsored by the Biological and Environmental Research program under Contract No. DE-AC05-76RL01830. Pacific Northwest National Laboratory is a multi-program national laboratory operated by Battelle for the DOE under Contract DE-AC05-76RL01830. Computational allocations from the 2024 ASCR Leadership Computing Challenge (ALCC) award.

We thank the computational resources from Tahoma (EMSL/PNNL), Deception (PNNL), Delta (NCSA/UIUC), and Polaris (ALCF).

Molecular graphics and analyses performed with UCSF ChimeraX, developed by the Resource for Biocomputing, Visualization, and Informatics at the University of California, San Francisco, with support from National Institutes of Health R01-GM129325 and the Office of Cyber Infrastructure and Computational Biology, National Institute of Allergy and Infectious Diseases. We thank Malio Nelson, Amity Andersen, and Scott Widmann for generating the homologue structural model of the dark complex for MED4.

## Author contributions

M.S.C., Z. L.S designed the research.

C.J., A.C., J.R., A.G., S.F. performed the computational research.

J.R., A.C., C.J., S.F., Z.L.S., and M.S.C. contributed to the analysis of the simulation data.

RW, S.F., W. S. annotated genomes

P.B., X.L., N.S., M.G., J.T., S.F., W.J.Q. performed experimental work to generate omics’ data

A.P., D.K., J.R., S.F., P.B., Z.J. contributed the analysis of the multimodal data for the computational research

A.P., J.E.E. performed experimental work to generate bioimaging data. A.G., D.K, A.C. segmented and transformed the tomograms for simulations.

L.A. packaged the multimodal data for release M.S.C., J.R. wrote the first draft of the paper. All authors contributed to writing the paper.

## Competing Interest Statement: No competing interest

Preprint Server:

## Classification

Physical Sciences/Biophysics and Computational Biology: Biological Sciences/ Systems Biology

## Notes

### Competing Interest Statement

The authors have declared no competing interest.

